# Cord blood DNA methylome in newborns later diagnosed with autism spectrum disorder reflects early dysregulation of neurodevelopmental and X-linked genes

**DOI:** 10.1101/850529

**Authors:** Charles E. Mordaunt, Julia M. Jianu, Ben Laufer, Yihui Zhu, Keith W. Dunaway, Kelly M. Bakulski, Jason I. Feinberg, Heather E. Volk, Kristen Lyall, Lisa A. Croen, Craig J. Newschaffer, Sally Ozonoff, Irva Hertz-Picciotto, M. Daniele Fallin, Rebecca J. Schmidt, Janine M. LaSalle

## Abstract

**Background:** Autism spectrum disorder (ASD) is a neurodevelopmental disorder with complex heritability and higher prevalence in males. Since the neonatal epigenome has the potential to reflect past interactions between genetic and environmental factors during early development, we performed whole-genome bisulfite sequencing of 152 umbilical cord blood samples from the MARBLES and EARLI high-familial risk prospective cohorts to identify an epigenomic signature of ASD at birth.

**Results:** We identified differentially-methylated regions (DMRs) stratified by sex that discriminated ASD from control cord blood samples in discovery and replication sets. At a region level, 7 DMRs in males and 31 DMRs in females replicated across two independent groups of subjects, while 537 DMR genes in males and 1762 DMR genes in females replicated by gene association. These DMR genes were significantly enriched for brain and embryonic expression, X chromosome location, and identification in prior epigenetic studies of ASD in post-mortem brain. In males and females, autosomal ASD DMRs were significantly enriched for promoter and bivalent chromatin states across most cell types, while sex differences were observed for X-linked ASD DMRs. Lastly, these DMRs identified in cord blood were significantly enriched for binding sites of methyl-sensitive transcription factors relevant to fetal brain development.

**Conclusions:** At birth, prior to the diagnosis of ASD, a distinct DNA methylation signature was detected in cord blood over regulatory regions and genes relevant to early fetal neurodevelopment. Differential cord methylation in ASD supports the developmental and sex-biased etiology of ASD, and provides novel insights for early diagnosis and therapy.

## Background

Autism spectrum disorder (ASD) is a heterogeneous neurodevelopmental disorder that affects 1 in 59 children in the United States [1]. ASD presents as persistent difficulties in social communication and interaction, restricted and repetitive behaviors and interests, as well as sensory sensitivities. Communication deficits can include delayed and monotonous speech, echolalia, poor verbal comprehension, difficulty understanding body language cues, and making eye contact, while behavioral deficits can include stereotyped movements, insistence on routine, fixated interests, and altered sensitivity to sensory input. ASD is currently diagnosed in childhood by one of several standardized scales that include interviews, behavioral observation, and clinical assessment, such as the gold standard Autism Diagnostic Observation Schedule (ADOS) [2, 3]. Clinical management of ASD symptoms consists of both behavioral interventions and pharmacological treatments. Intensive behavioral interventions have been associated with better outcomes in children with ASD, especially for those diagnosed within the first 2-3 years of life as they are the most amenable to behavioral intervention [4].

Risk for ASD is thought to originate from a combination of multiple genetic and environmental factors, as well as gene-environment interactions [5]. The genetic architecture of ASD includes both rare variants with strong effects and multiple common variants with weak individual effects [6]. Genes associated with ASD in genetic studies are enriched for pathways affecting neuronal homeostasis and embryonic development and include synaptic neurotransmitter receptors, cytoskeletal proteins, protein degradation factors, and chromatin regulators. However, in keeping with the complexity of ASD, no single genetic variant has been found that accounts for more than 1% of cases. Environmental factors are also known to contribute to ASD risk, especially during the prenatal period [7, 8]. During gestation, when rapid cellular proliferation and differentiation is occurring, environmental stressors can have long-lasting effects on behavior [9]. Maternal factors shown to modify ASD risk include environmental toxicants, nutritional factors, fever, pre-eclampsia, gestational diabetes, and medications [10]. ASD etiology is thought to begin early in development through interactions between genetic and environmental factors that alter neurodevelopmental trajectories [11, 12].

ASD has consistently been found to be more frequent in males compared to females, at about a 3 to 1 ratio [13]. In autistic individuals without intellectual disability, the ratio is increased to 11 to 1 [14]. The sex ratio has been explained in two ways, which are not mutually exclusive: 1) differential ascertainment, where females tend to show more social motivation and ability to hide their social difficulties; and 2) a female protective effect, where females with the same symptoms as males have a larger burden of genetic mutations. The recurrence rate for both males and females in families containing a female proband is higher than those with only male probands, suggesting a higher genetic load [15, 16]. Females diagnosed with ASD may also have more severe symptoms and comorbidities than their male counterparts [17]. A female protective effect in ASD etiology may be explained by the presence of two copies of chromosome X, which has a disproportionate number of genes involved in neurodevelopment [14]. Through the process of X chromosome inactivation, females are mosaics for mutations in X-linked genes, resulting in a diluted effect of harmful X-linked mutations. Interestingly, some genes on the X chromosome without Y-linked homologs escape X chromosome inactivation, resulting in higher expression in females than in males, and potentially impacting sex-specific susceptibility to ASD [18].

Epigenetic mechanisms have been proposed to account for the sex bias, gene-environment interactions, and developmental origins of ASD etiology [19]. Epigenetic modifications, including DNA methylation, histone post-translational modifications (PTMs), noncoding RNA, and chromatin architecture, are influenced by both genetic and environmental factors and are established dynamically during development to reinforce cell lineage and function [20]. DNA methylation, which primarily occurs as the addition of a methyl group to the 5^th^ carbon of the cytosine in a CpG dinucleotide, is the best understood epigenetic modification and the most tractable for large human studies. Epigenome-wide association studies (EWAS) have identified locus-specific differential methylation and epigenetic signatures associated with ASD. EWAS in post-mortem brain have identified differential methylation of genes involved in synaptic transmission and the immune response, with a particular enrichment for open chromatin regions and genes important for microglia [21–24]. Similarities in brain epigenetic dysregulation have been found between individuals with idiopathic and syndromic ASDs, including Dup15q and Rett syndrome [23,25,26]. Tissues accessible in humans, such as placenta, paternal sperm, buccal, and blood, have been used to identify differential methylation in ASD by EWAS [27–31], but few individual loci have been replicated. Previous EWAS have been hampered by study design limitations, including case-control cohorts that may be confounded by reverse causation and microarray-based platforms that cover less than 3% of the CpG sites in the genome [32].

Here, we obtained umbilical cord blood samples from ASD and typically developing (TD) subjects from two high-familial risk prospective cohorts (i.e., cohorts following younger siblings of a child already diagnosed with ASD through that subsequent child’s early development) in order to use an accessible tissue at birth before ASD symptoms developed. We used whole-genome bisulfite sequencing (WGBS) to assess levels of DNA methylation across more than 20 million CpG sites and sex-stratified the analysis to reveal epigenetic differences specific to males or females. Sex-specific differentially-methylated regions (DMRs) distinguishing ASD from TD newborns were analyzed with a systems biology-based approach to identify impacted genes, pathways, transcription factors, and chromatin states. Unlike most previous ASD EWAS, this approach examines methylation before the onset of symptoms, correlates newborn methylation with quantitative clinical measurements at 36 months, covers the majority of CpG sites in the genome, and emphasizes functional genomic regions. As a result of using this approach, we identified sex-specific epigenomic signatures of ASD in umbilical cord blood that replicated in an independent group of subjects.

## Results

### Study subject characteristics in relation to neurodevelopmental outcome at 36 months

Subjects in this study were from the Markers of Autism Risk in Babies - Learning Early Signs (MARBLES; TD *n* = 44, ASD *n* = 44) and Early Autism Risk Longitudinal Investigation (EARLI; TD *n* = 32, ASD *n* = 32) high-familial risk prospective cohorts, and they include those with an ASD outcome and those determined to be TD at 36 months. By definition of the high-familial risk cohort design, all subjects have an older sibling with ASD. Additionally, due to the sex bias in ASD, the majority (74%) of the subjects are males (Additional file 2: Table S1, S2). The ADOS assessment scale is a criteria in the diagnostic algorithm for ASD (see Methods), and as such the ASD subjects exhibit significantly higher severity on the ADOS comparison score [3] (ASD mean = 6.6, TD mean = 1.1, *q* = 8.0E-62). The Mullen Scales of Early Learning (MSEL) [33] measures cognitive functioning and was used to exclude subjects with intellectual disability from those classified with TD; thus ASD subjects also had lower performance on the MSEL early learning composite score (ASD mean = 80, TD mean = 112, *q* = 4.2E-21) when compared to TD subjects. Mothers of ASD subjects differed from mothers of TD subjects on a few characteristics. Specifically, mothers of ASD subjects had altered educational attainment (*p* = 0.02) including increased attainment of at most an associate’s degree (ASD = 24%, TD = 11%) and decreased attainment of at most a bachelor’s degree (ASD = 20%, TD = 43%). They also were less likely to own their home (ASD = 50%, TD = 70%, *p* = 0.01), had a higher pre-pregnancy body mass index (ASD mean = 29, TD mean = 27, *p* = 0.04), and were more likely to smoke during pregnancy (ASD = 11%, TD = 1%, *p* = 0.03).

**Table 1.** ASD DMR genes replicate in males and females in independent sample sets.

| Gene Set 1 | Genes | Gene Set 2 | Genes | Overlap | % Overlap | Odds Ratio | *p*-value |
| --- | --- | --- | --- | --- | --- | --- | --- |
| Discovery Males<br>(*n* = 74) | 949 | Replication Males<br>(*n* = 38) | 5429 | 537 | 56.6 | 5.13 | **1.2E-126** |
| Discovery Females<br>(*n* = 32) | 2912 | Replication Females<br>(*n* = 8) | 9496 | 1762 | 60.5 | 2.86 | **3.6E-153** |
| Replicated Males<br>(All chrs) | 537 | Replicated Females<br>(All chrs) | 1762 | 162 | 30.2 | 6.20 | **4.0E-60** |
| Replicated Males<br>(chrX) | 55 | Replicated Females<br>(chrX) | 173 | 21 | 38.2 | 3.48 | **4.6E-5** |

### Lower global DNA methylation in umbilical cord blood from males later diagnosed with ASD corresponds with increased nucleated red blood cells

ASD subjects were found to have lower global CpG methylation than TD subjects (Additional file 2: Table S1). Because this has been found previously in blood from children with ASD [34], we investigated this association in more detail. Specifically, analyses were performed stratified by sex, both within and pooled across sequencing platforms, and included adjustment for duplicate reads and sequencing platform. Only males with ASD were significantly hypomethylated compared to males with TD (pooled estimate = −0.55% methylation, *q* = 0.02), while females with ASD had similar global methylation as females with TD (pooled estimate = +0.14% methylation, *q* = 0.92; Additional file 1: Fig. S1A, S2A, Additional file 2: Table S3). Furthermore, scores on the MSEL were positively associated with global methylation only in males, where male subjects with lower scores also had lower global methylation (pooled estimate = +0.30% methylation per standard deviation (SD), *q* = 0.02; Additional file 1: Fig. S1). Other examined demographic and technical covariates were not associated with global methylation, with the exception of the proportion of nucleated red blood cells (nRBCs) estimated based on percent methylation at cell-type-specific loci [35] (Additional file 1: Fig. S1A, S2D, Additional File 2: Table S3). In both males and females, nRBCs were negatively associated with global methylation (pooled males estimate = −0.71% methylation per SD, *q* = 8.6E-18; pooled females estimate = −0.71% methylation per SD, *q* = 7.0E-4). Similar to the pattern observed with global methylation, nRBCs were increased only in males with ASD compared to males with TD (pooled estimate = +2.65% nRBCs, *q* = 0.046; Additional file 1: Fig. S2B, Additional file 2: Table S3), and were also negatively associated with MSEL scores only in males (pooled estimate = −1.21% nRBCs per SD, *q* = 0.046; Additional file 1: Fig. S2C). Importantly, when nRBCs were added as an adjustment covariate, global methylation was no longer associated with ASD diagnosis in males (pooled estimate = −0.18% methylation, *p* = 0.19; Additional file 2: Table S3). These findings suggest that an increased proportion of nRBCs in whole cord blood, specifically in males later diagnosed with ASD, can explain their lower observed global methylation levels.

### Region-specific differential methylation patterns in umbilical cord blood distinguish males and females later diagnosed with ASD from TD controls

Because DNA methylation, like ASD etiology, is influenced by both genetic and environmental factors during prenatal life, we hypothesized that umbilical cord blood DNA from newborns who later develop ASD would exhibit DMRs compared to those with TD. To test this hypothesis, we analyzed cord blood WGBS data from the MARBLES and EARLI studies for DMRs with methylation differences of more than 5% between ASD and TD samples and permutation-based empirical *p*-values less than 0.05. These data were all generated on the HiSeq X Ten sequencing platform and represent the “discovery set” (males TD *n* = 39, ASD *n* = 35; females TD *n* = 17, ASD *n* = 15). Because of the expected sex differences in DNA methylation patterns due to X chromosome inactivation and the previously observed sex differences in subjects with ASD [14], we performed a sex-stratified analysis to preserve sex-specific differential methylation. In males, 150 hypermethylated (ASD greater than TD) and 485 hypomethylated (ASD less than TD) DMRs associated with ASD were identified, and methylation in these regions clustered subjects distinctly by outcome (Fig. 1A-B, Additional file 2: Table S4). Similarly, ASD DMRs were identified in females, including 863 hypermethylated and 1089 hypomethylated DMRs that could distinguish between subjects with ASD and TD outcome (Fig. 1C-D, Additional file 2: Table S5). When methylation levels within male or female ASD DMRs were compared with subject characteristics, they were specifically associated with autism severity and MSEL cognitive scores but not other demographic and technical variables (Additional file 1: Fig. S3, Additional file 2: Table S4, S5). Notably, the majority of ASD DMRs were not associated with the proportion of nRBCs, which was a driver of global methylation (*p* > 0.05, males: 77% of DMRs not associated, females: 94% of DMRs not associated). An alternate approach of combining males and females, with sex included as an adjustment covariate, revealed similar results but fewer ASD-associated DMRs (Additional file 1: Fig. S4, Additional file 2: Table S6).

**Figure 1.**
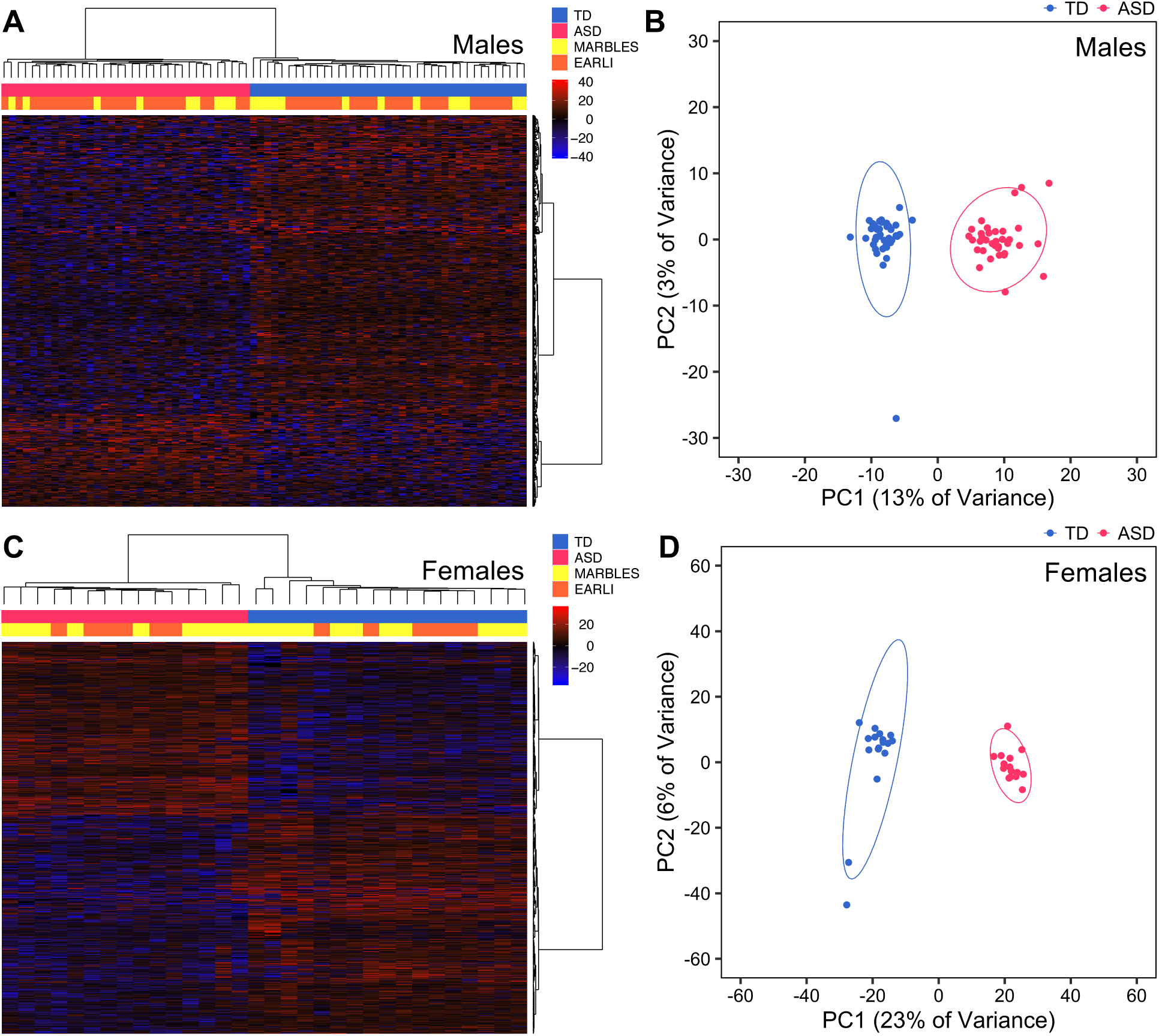
DMRs identified in males and females distinguish ASD from TD subjects in the discovery set. (A) Heatmap or (B) principal component analysis (PCA) plot using percent methylation for each sample in autism spectrum disorder (ASD) versus typically developing (TD) differentially-methylated regions (DMRs) identified in cord blood from male discovery subjects (150 hypermethylated DMRs, 485 hypomethylated DMRs; TD *n* = 39, ASD *n* = 35). (C) Heatmap or (D) PCA plot using percent methylation for each sample at ASD DMRs identified in cord blood from female discovery subjects (863 hypermethylated DMRs, 1089 hypomethylated DMRs; TD *n* = 17, ASD *n* = 15). For heatmaps, subjects are colored by diagnostic group and study, and methylation is relative to the mean for each DMR. For PCA plots, ellipses indicate 95% confidence limits. EARLI, Early Autism Risk Longitudinal Investigation; MARBLES, Markers of Autism Risk in Babies -Learning Early Signs;

To validate our finding that DMRs between ASD and TD subjects are present in cord blood, we replicated the WGBS on an independent group of subjects in the MARBLES study using a different sequencing platform. Data from these subjects were all generated on the HiSeq 4000 sequencing platform and represent the “replication set” (males TD *n* = 17, ASD *n* = 21; females TD *n* = 3, ASD *n* = 5). Similar to the discovery set, sex-stratified DMRs were identified in the replication set that could specifically cluster ASD versus TD subjects and were more strongly associated with autism severity and cognition than other variables (Additional file 1: Fig. S5, S6, Additional file 2: Table S7, S8). Again, most ASD DMRs in males or females were not associated with the proportion of nRBCs (*p* > 0.05, males: 75% of DMRs not associated, females: 98% of DMRs not associated). DMRs were also identified in the replication set with a combined sex-adjusted approach, which similarly resulted in fewer DMRs than the sex-stratified analysis (Additional file 1: Fig. S7, Additional file 2: Table S9). To computationally validate the detected DMRs, we redid all comparisons with a different statistical approach, as implemented in the bsseq R package [36]. DMRs in all comparisons significantly overlapped between the two methods, suggesting these ASD DMRs identified in cord blood are not an artifact of the computational approach (permutation test, z-score > 35 and *p* < 1.0E-4 for all; Additional file 2: Table S10). Interestingly, when we compared the ability to replicate the WGBS-derived ASD DMRs with Infinium HumanMethylation450 or MethylationEPIC arrays, 82% and 84% of DMRs from the male discovery and replication sets, respectively, did not overlap even one array probe (Additional file 1: Fig. S8). Similarly, 71% and 69% of DMRs identified in females from the discovery and replication sets, respectively, were not covered on either array, reinforcing the utility of the WGBS approach. Together, these results demonstrate the discovery and technical replication of novel DMRs in cord blood associated with ASD outcome at 36 months in two high-familial risk cohorts.

### ASD DMRs in umbilical cord blood replicate across independent groups of subjects

After confirming that we could identify ASD DMRs in cord blood, we next hypothesized that specific regions are consistently differentially methylated in both discovery and replication sets of newborns later diagnosed with ASD. Testing for the replication of individual DMRs was undertaken with three approaches: region overlap, differential methylation, and gene overlap. When DMRs identified in males from the discovery set were overlapped by genomic location with those found in males from the replication set, 4 hypermethylated DMRs and 3 hypomethylated DMRs were present in both (out of 635 discovery DMRs), which was more than expected by chance (permutation test, hypermethylated z-score = 8.1, *p* < 1.0E-4; hypomethylated z-score = 6.1, *p* < 1.0E-4; Additional file 2: Table S11, S12). In females, 7 hypermethylated and 24 hypomethylated DMRs were identified in both subject sets (out of 1954 discovery DMRs), also a significant overlap (permutation test, hypermethylated z-score = 14.2, *p* < 1.0E-4; hypomethylated z-score = 23.7, *p* < 1.0E-4). To determine whether ASD DMRs detected in the discovery set showed the same directional differential methylation in the replication set, we compared the percent methylation over each of these regions to ASD outcome in both sample sets. In males, 15 out of 635 DMRs in the discovery set were differentially methylated similarly in the replication set, while 23 out of 1954 discovery DMRs were directionally similar in females from both sample sets (*p* < 0.05; Additional file 2: Table S13). We also compared methylation at diagnosis DMRs from the discovery set to symptom severity within ASD subjects and identified 1 out of 635 DMRs in males and 3 out of 1954 DMRs in females with consistent correlation in both sample sets (*p* < 0.05; Additional file 1: Fig. S9, Additional file 2: Table S14).

Because the functional output of the genome is the transcription of genes, we compared genes annotated to ASD DMRs in each subject set. For males, a significant enrichment of 537 out of 949 DMR genes in the discovery set were also in the replication set (odds ratio = 5.1, *p* = 1.2E-126; Table 1, Additional file 2: Table S15). Similarly, for females, DMR genes in the discovery set significantly overlapped with those in the replication set, with 1762 out of 2912 present in both (odds ratio = 2.9, *p* = 3.6E-153). Replicated DMR genes in males also significantly overlapped with those in females, with 162 genes replicated in both sexes (odds ratio = 6.2, *p* = 4.0E-60). However, the majority of replicated ASD DMR genes were specific to males or females (375/537 male-specific, 1600/1762 female-specific). DMR genes also replicated in the combined sex-adjusted analysis, but most of the genes identified when stratifying for sex were unique (odds ratio = 5.5, *p* = 1.9E-45; Additional file 1: Fig. S10). These findings suggest that reproducible ASD DMRs are present in cord blood, and these are not dependent on the particular group of subjects or technical approaches.

### Genes in ASD differentially-methylated blocks (DMBs) replicate between independent groups of subjects and are enriched for ASD DMR genes, cadherins, and developmental genes

Because of the differences detected in global CpG methylation, we also tested the hypothesis that large DMBs (defined as regions more than 5 kilobases in length with greater than 1% methylation difference in ASD versus TD) are present in cord blood DNA from individuals with ASD. DMBs were indeed present in males and females from both the discovery and replication sets (Additional file 2: Table S16-S19). A comparison of genes annotated to DMBs in the discovery and replication sets revealed a significant overlap of 58 genes in males and 23 genes in females that were present across both sets (Males odds ratio = 3.7, *p* = 1.7E-13; Females odds ratio = 2.6, *p* = 1.7E-4; Additional file 2: Table S15, S20). A significant enrichment of replicated cord blood ASD DMR genes was observed in these replicated DMB genes, including 20 genes in males and 12 genes in females (Males odds ratio = 25.8, *p* = 1.3E-19; Females odds ratio = 15.1, *p* = 6.1E-9; Additional file 1: Fig. S11, Additional file 2: Table S15, S20). In males and females from both subject sets, DMB genes were significantly enriched for chromosome 5 location, cell adhesion and calcium signaling functions, and expression during embryonic development (*q* < 0.05; Additional file 1: Fig. S12, Additional file 3: Table S21). Male DMB genes were specifically enriched for expression in brain and bone marrow endothelial cells, while female DMB genes were especially enriched for cadherin genes (*q* < 0.05). These findings suggest that ASD DMBs are present in cord blood, and they impact some of the same genes and pathways as ASD DMRs, with a particular impact on cadherins.

### Cord blood ASD DMR genes are enriched for neurodevelopmental genes on the X chromosome that are epigenetically dysregulated in ASD brain

To test the hypothesis that ASD DMR genes identified in cord blood are functionally relevant to ASD etiology, we examined these genes for enrichment in predefined gene sets including pathways, chromosomal location, tissue expression, and previous genome-wide studies of ASD. For both the discovery and replication sets, ASD DMR genes detected in males and females were significantly enriched for X chromosome location and functions localized to the postsynaptic membrane (*q* < 0.05; Fig. 2, Additional file 3: Table S22). ASD DMR genes identified from male and female cord blood were also enriched for expression in embryonic development, multiple brain regions, pituitary, and testes (*q* < 0.05). Genes that replicated in both males and females and are expressed in brain, at the postsynaptic membrane, and during embryonic development include *GABRA2*, *GABRG1*, *GRIA3*, *GRIK2*, *LRRC4C*, *LRRTM1*, *LRRTM4*, *KCNC2*, and *ZC4H2*. In males but not females, cord blood ASD DMR genes were enriched for locations on chromosomes 4 and 8, expression in cingulate cortex, temporal lobe, and blood, and for functions in neuroactive ligand-receptor interactions (*q* < 0.05). In contrast, female DMR genes identified in cord blood were enriched for genes associated with chromosome 18, mental retardation, dendritic spines, and calcium, as well as dorsal root ganglia and subthalamic nucleus expression (*q* < 0.05). ASD DMR genes in cord blood were enriched for X chromosome genes in both males and females and these significantly overlapped, with 21 genes in common (odds ratio = 3.5, *p* = 4.6E-5; Table 1, Fig. 3). However, most replicated genes on the X chromosome were specific for either males or females, with more DMR genes detected in females (34/55 male-specific, 152/173 female-specific). Interestingly, all genes on the X chromosome (not just DMR genes) are enriched for brain and embryonic expression, as well as mental retardation and autism, compared to all genes in the genome (*q* < 0.05; Additional file 1: Fig. S13, Additional file 3: Table S23). Accordingly, many replicated X-linked ASD DMR genes are expressed in fetal brain (Additional file 1: Fig. S14, S15).

**Figure 2.**
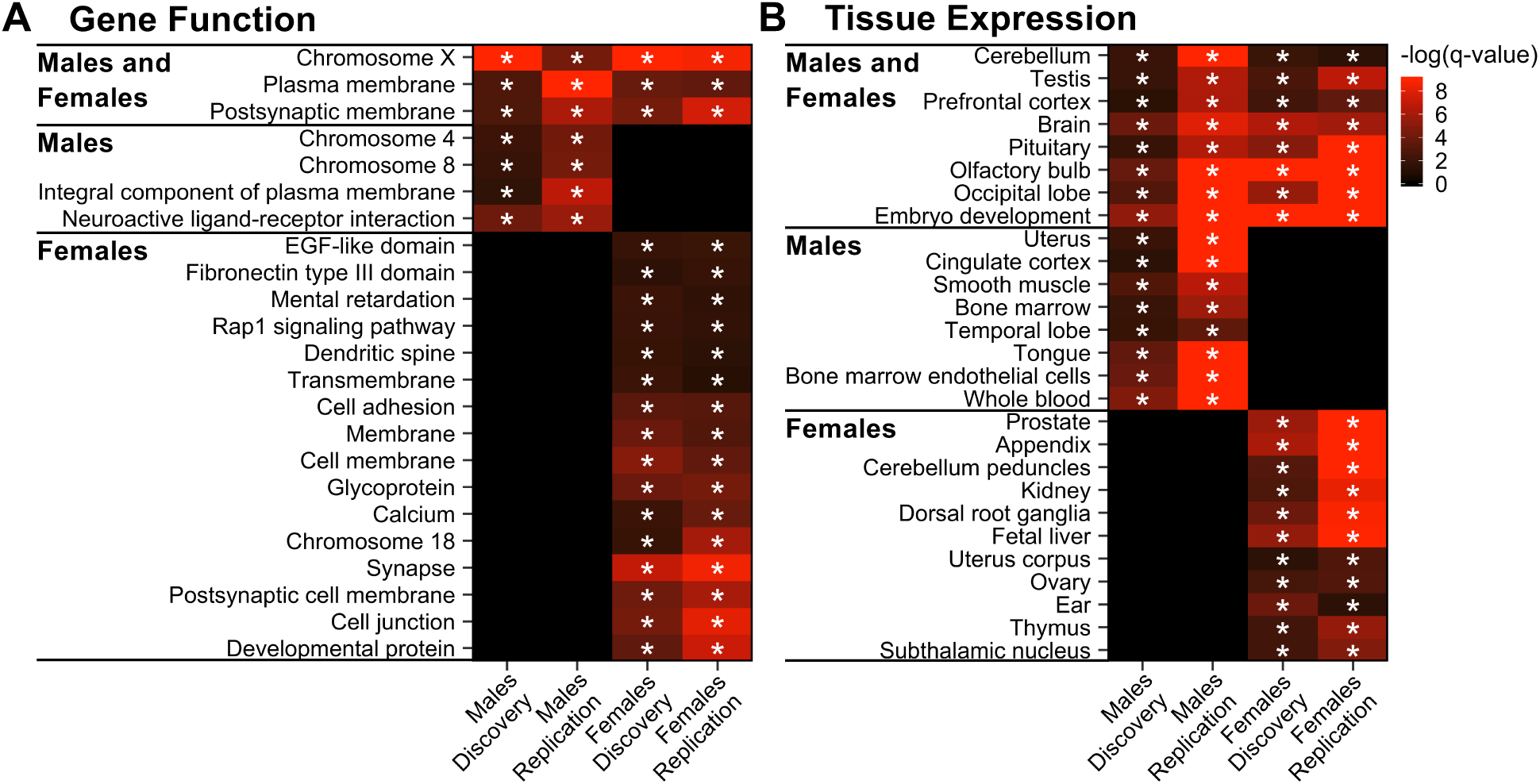
ASD DMR genes are significantly enriched for X-linked and synaptic genes in males and females. Terms significantly enriched among cord blood ASD DMR genes in both discovery and replication sample sets for either males or females (* *q* < 0.05). Heatmaps show - log_10_(*q*-value) for enrichment in genes annotated to DMRs relative to genes annotated to background calculated using the Database for Annotation, Visualization, and Integrated Discovery (DAVID) for (A) all categories except tissue expression or (B) tissue expression categories only. Both heatmaps were plotted using the same scale and terms were sorted by replicated sex (pooled males TD *n* = 56, ASD *n* = 56; pooled females TD *n* = 20, ASD *n* = 20).

**Figure 3.**
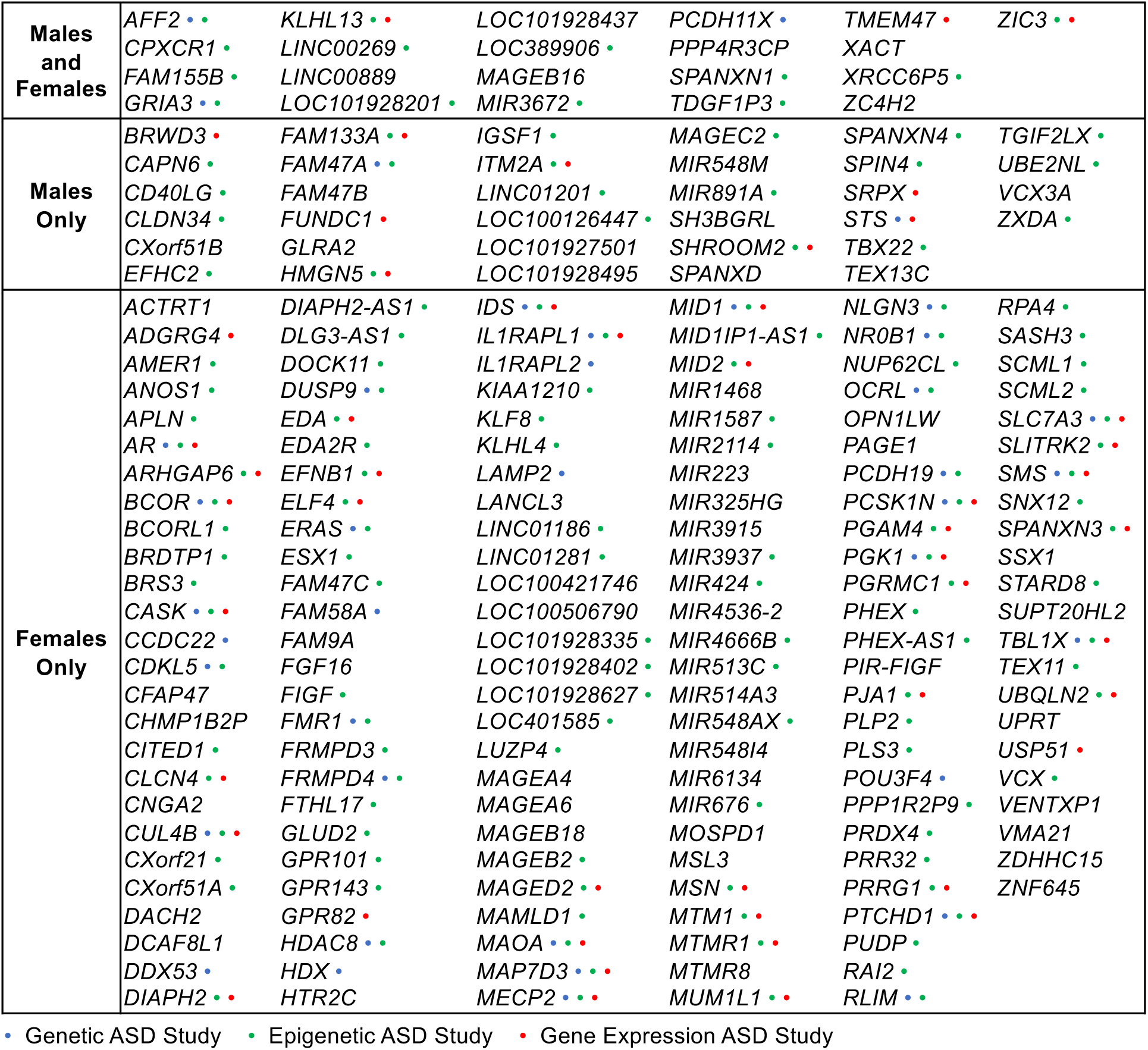
Replicated sex-independent, male-specific, and female-specific X-linked DMR genes and their overlap with previous ASD studies. Genes on the X chromosome that were annotated to cord blood ASD DMR genes in both discovery and replication sample sets for males and/or females. Dots indicate overlap with at least 1 genetic, epigenetic, or gene expression study of ASD.

To further examine relevance to ASD etiology, we overlapped cord blood ASD DMR genes with previously reported gene sets from genetic, epigenetic, and transcriptional studies of ASD samples. The majority (77%) of the replicated X chromosome cord blood ASD DMR genes in males or females were identified in previous studies of ASD (Fig. 3). Genome-wide, cord blood ASD DMR genes in males and females from both the discovery and replication sets were significantly enriched for genes identified in previous studies examining DNA methylation (by both array-based and WGBS approaches) in the cingulate cortex, prefrontal cortex, and temporal cortex, and also histone 3 lysine 27 (H3K27) acetylation in prefrontal cortex and cerebellum from subjects with ASD (*q* < 0.05; Fig. 4 shows replicated enrichments, Additional file 1: Fig. S16 shows all comparisons, Additional file 3: Table S24 shows statistics, genes, and sources). Notably, 16 genes that replicated in both males and females from our WGBS analyses were also identified in at least four epigenetic studies of ASD post-mortem brain, including *CHST15*, *CPXM2*, *FAM49A*, *FAM155B*, *KDR*, *LINC01491*, *LINC01515*, *LOC100506585*, *LOC100996664*, *LOC101928441*, *MIR378C*, *RBFOX1*, *RBMS3-AS1*, *ST6GAL2*, *TGFBR2*, and *TIMP3*. Many of the genes also identified in previous epigenomic studies of ASD are located on the X chromosome and expressed in fetal brain, including *GRIA3*, *AFF2*, and *KLHL13*, which replicated in males and females; *SHROOM2*, *ZXDA*, and *HMGN5*, which were male-specific; and *MECP2*, *FMR1*, and *PCSK1N*, which were female-specific (Additional file 1: Fig. S17-S19). Male and female DMR genes also significantly overlapped those identified as differentially methylated in prefrontal cortex from the syndromic ASDs Dup15q and Rett syndromes, and in sperm from fathers of toddlers displaying ASD-like traits in EARLI (*q* < 0.05). Female but not male ASD DMR genes were significantly enriched for dysregulated gene sets from other ASD DNA methylation studies in brain, but also in MARBLES placenta, lymphoblast cell lines, and newborn blood spots (*q* < 0.05). Additionally, female ASD DMR genes significantly overlapped with known ASD risk genes and those associated with altered histone 3 lysine 4 (H3K4) trimethylation in neurons from ASD subjects (*q* < 0.05).

**Figure 4.**
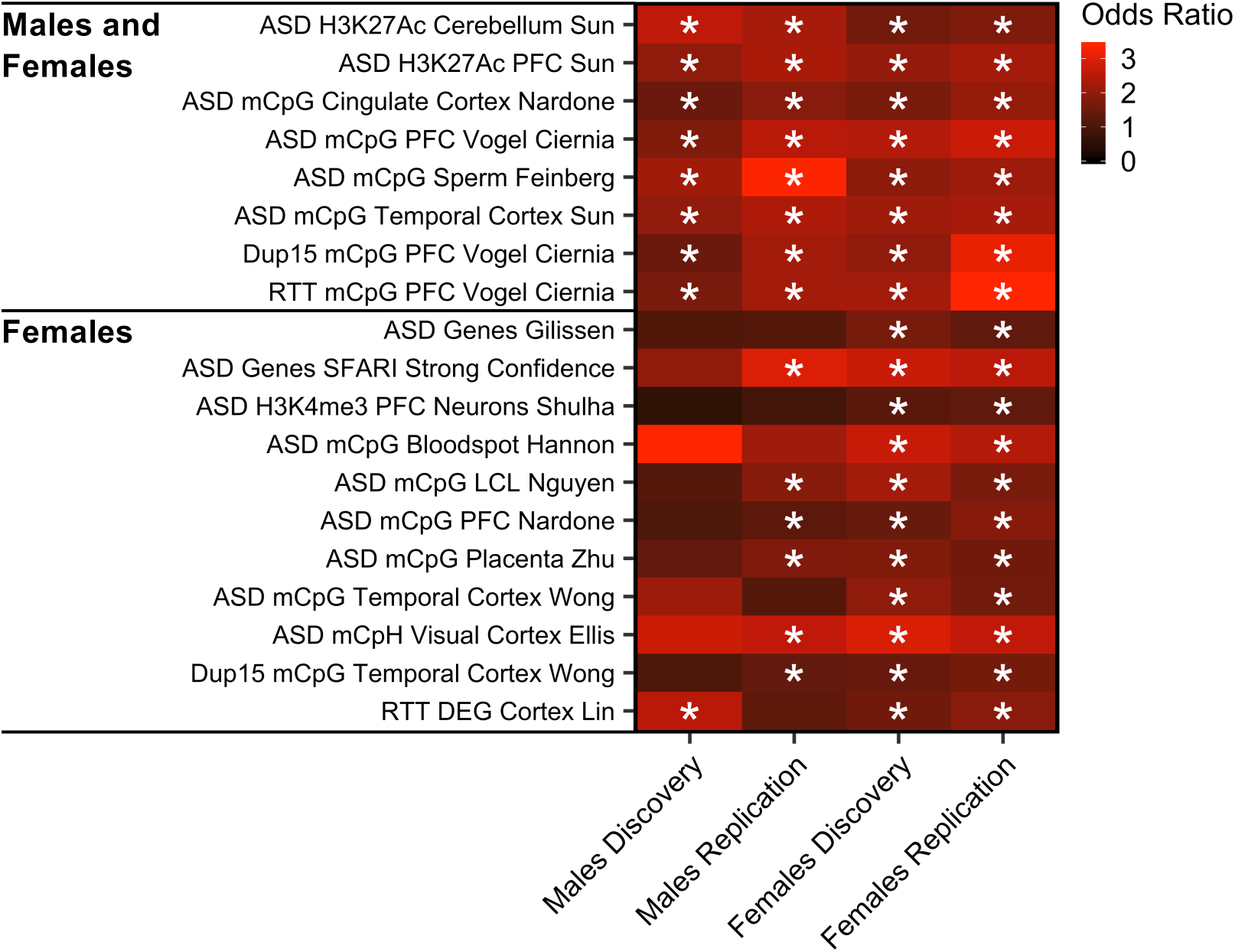
Cord blood ASD DMR genes are significantly enriched for epigenetically dysregulated genes in ASD brain. Gene sets significantly enriched among ASD DMR genes in both discovery and replication sample sets for either males or females (* *q* < 0.05). Heatmaps show odds ratios for enrichment in genes annotated to DMRs relative to genes annotated to background calculated with Fisher’s exact test for previously published studies of ASD and other neurological disorders. *P*-values were adjusted using the false discovery rate (FDR) method for the total number of gene lists compared (pooled males TD *n* = 56, ASD *n* = 56; pooled females TD *n* = 20, ASD *n* = 20). Ac, acetylation; DEG, differentially-expressed genes; Dup15, Chromosome 15q11-q13 Duplication syndrome; H3K27, histone 3 lysine 27; H3K4, histone 3 lysine 4; LCL, lymphoblastoid cell line; mCpG, CpG methylation; mCpH, CpH methylation; me3, trimethylation; PFC, prefrontal cortex; RTT, Rett syndrome; SFARI, Simons Foundation Autism Research Initiative;

In striking contrast to the strong concordance across epigenetic ASD studies using different marks and methodologies, no study examining differential gene expression in ASD subjects showed a significant overlap in gene hits with any of our ASD DMR genes from the discovery or replication set, including one conducted in cord blood from the MARBLES and EARLI cohorts [37] (Additional file 1: Fig. S16, Additional file 3: Table S24). The extensive overlap of cord blood DMR genes with those identified in other epigenomic studies across several tissues, combined with the lack of overlap with differentially expressed genes, supports the hypothesis that the DNA methylome is less time dependent than the transcriptome and more likely reflects past alterations that occurred in early development.

### ASD DMRs are enriched for a pan-tissue epigenomic signature that differs between males and females on the X chromosome

To test the hypothesis that cord blood ASD DMRs were reflecting chromatin differences in *cis*-regulatory regions important in early prenatal life, we examined the enrichment of autosomal and X chromosomal DMRs for histone PTMs and chromatin states across many tissues and stages. Cord blood ASD DMRs were positionally compared with histone PTM chromatin immunoprecipitation sequencing (ChIP-seq) peaks and ChromHMM-defined chromatin states in 127 cell types from the Roadmap Epigenomics Project [30]. ASD hyper- and hypomethylated DMRs in males and females in both sample sets were enriched for chromatin states involved in transcriptional regulation, including active transcription start sites (TssA and TssAFlnk) and bivalent enhancers (EnhBiv), across tissue types (*q* < 0.05; Fig. 5A,C; Additional file 1: Fig. S20A,C, Additional file 3: Table S25-S28). These chromatin states corresponded to significant enrichment in regions with H3K4me3, H3K27me3, and H3K4me1 across tissues (*q* < 0.05; Additional file 1: Fig. S21A,C, S22A,C, Additional File 3: Table S25-S28). In males, hypermethylated DMRs were enriched in genic enhancers (EnhG), while hypomethylated DMRs were enriched near bivalent (BivFlnk) and polycomb-repressed (ReprPC) regions in both sample sets (*q* < 0.05). In females, both hyper- and hypomethylated DMRs were enriched in EnhG, BivFlnk, and ReprPC regions in discovery and replication sets (*q* < 0.05). Together, these results demonstrate a sex-independent epigenetic signature of poised bivalent genes and enhancers that is pan-tissue, rather than limited to blood or immune-specific functions.

**Figure 5.**
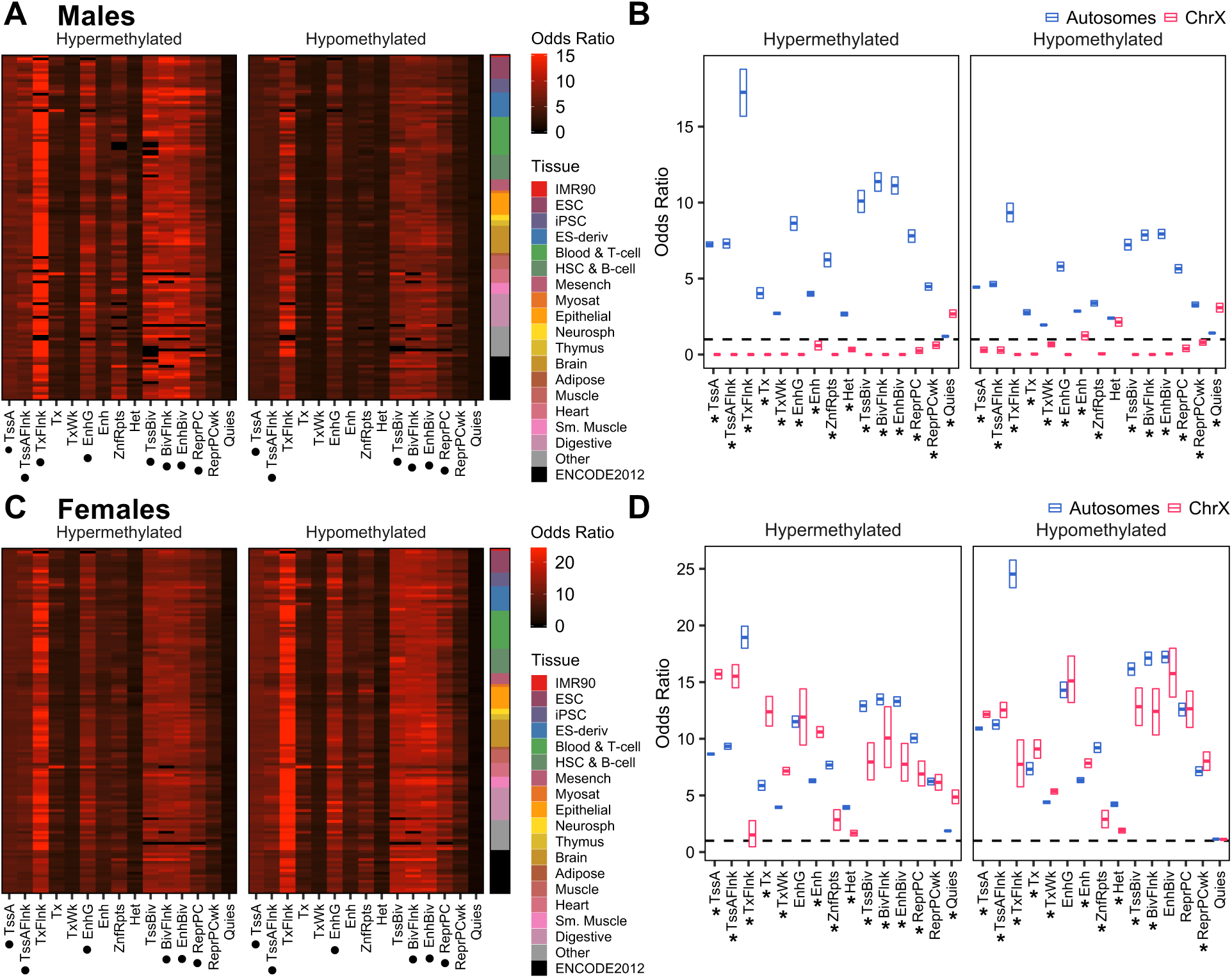
Pan-tissue chromatin signature of cord blood ASD DMRs reveals X-linked sex differences in chromatin states. Cord blood ASD DMRs from the discovery set were overlapped with 15-state model ChromHMM segmentations from 127 cell types in the Roadmap Epigenomics Project using the Locus Overlap Analysis (LOLA) R package. (A, C) The enrichment odds ratio was plotted for hypermethylated and hypomethylated DMRs identified in (A) males or (C) females from the discovery set. Top enriched (•) chromatin states were identified as those with odds ratio and -log(*q*-value) of at least the median value for that DMR set and with a *q*-value less than 0.05 for more than half of all cell types. (B, D) The enrichment odds ratio was plotted for hypermethylated and hypomethylated DMRs on autosomes or the X chromosome identified in (B) males or (D) females from the discovery set. Boxes represent mean and 95% confidence limits by nonparametric bootstrapping. Significance of differential enrichment of X chromosome compared to autosome DMRs was assessed by paired t-test of odds ratios for each cell type. *P*-values were adjusted for the number of chromatin states using the FDR method (* *q* < 0.05, males TD *n* = 39, ASD *n* = 35; females TD *n* = 17, ASD *n* = 15). BivFlnk, flanking bivalent transcription start site or enhancer; Enh, enhancer; EnhBiv, bivalent enhancer; EnhG, genic enhancer; Het, heterochromatin; Quies, quiescent region; ReprPC, polycomb-repressed region; ReprPCWk, weak polycomb-repressed region; TssA, active transcription start site; TssAFlnk, flanking active transcription start site; TssBiv, bivalent transcription start site; Tx, strong transcription; TxFlnk, transcribed at gene 5’ and 3’; TxWk, weak transcription; ZnfRpts, zinc finger genes and repeats;

Interestingly, males and females displayed divergent patterns of ASD DMR enrichment in chromatin states and histone PTMs on the X chromosome versus autosomes. In males, hyper- and hypomethylated X chromosomal ASD DMRs in both sample sets were uniquely enriched for quiescent regions (Quies), and less enriched for all other chromatin states and corresponding histone PTMs compared to autosomal DMRs (*q* < 0.05; Fig. 5B, Additional file 1: Fig. S20B, S21B, S22B, Additional file 3: Table S25, S27). In contrast to males, X-linked ASD DMRs in females were strikingly more enriched in active transcribed chromatin states (TssA, TssAFlnk, Tx, TxWk, and Enh) but less enriched in flanking transcribed regions (TxFlnk) and heterochromatin (Het) compared to autosomal DMRs (*q* < 0.05; Fig. 5D, Additional file 1: Fig. S20D, Additional file 3: Table S26, S28). Hypermethylated ASD DMRs in females were also less enriched in other repressed or bivalent regions (ZnfRpts, TssBiv, BivFlnk, EnhBiv, and ReprRC) compared to autosomal DMRs (*q* < 0.05). These chromatin state differences in female X-linked DMRs corresponded with an increased enrichment for H3K4me1, H3K4me3, and H3K36me3 and a decreased enrichment for H3K9me3 (*q* < 0.05, Additional file 1: Fig. S21D, S22D). A CpG island analysis also confirmed the chromatin state enrichment differences between autosomal and X-linked ASD DMRs in males versus females (Additional file 1: Fig. S23, Additional file 3: Table S29). Together, these results suggest that in both males and females, autosomal ASD DMRs represent euchromatic, actively repressed, or bivalent chromatin states; while male-specific X-linked ASD DMRs reflect repressed quiescent heterochromatic regions; and female-specific X-linked ASD DMRs encompass euchromatic, transcriptionally-active regions.

### ASD DMRs are enriched for binding motifs of methylation-sensitive transcription factors relevant to the fetal brain epigenome

Since DNA methylation may modify the actions of transcription factors, we hypothesized that specific methyl-sensitive transcription factor binding sites could be identified in ASD DMRs, which may give further insight into their functional relevance in early development and X-linked sex differences. To test this hypothesis, we examined enrichments of transcription factor binding site motifs within sex-specific ASD DMRs split by autosome versus X chromosome location. In both males and females, hypomethylated autosomal DMRs were enriched in motifs for HOXA2, TBX20, and ZNF675 (*q* < 0.05; Fig. 6A, Additional file 3: Table S30). However, for X-linked DMRs, none of the top enriched motifs were in common between males and females (Fig. 6B).

**Figure 6.**
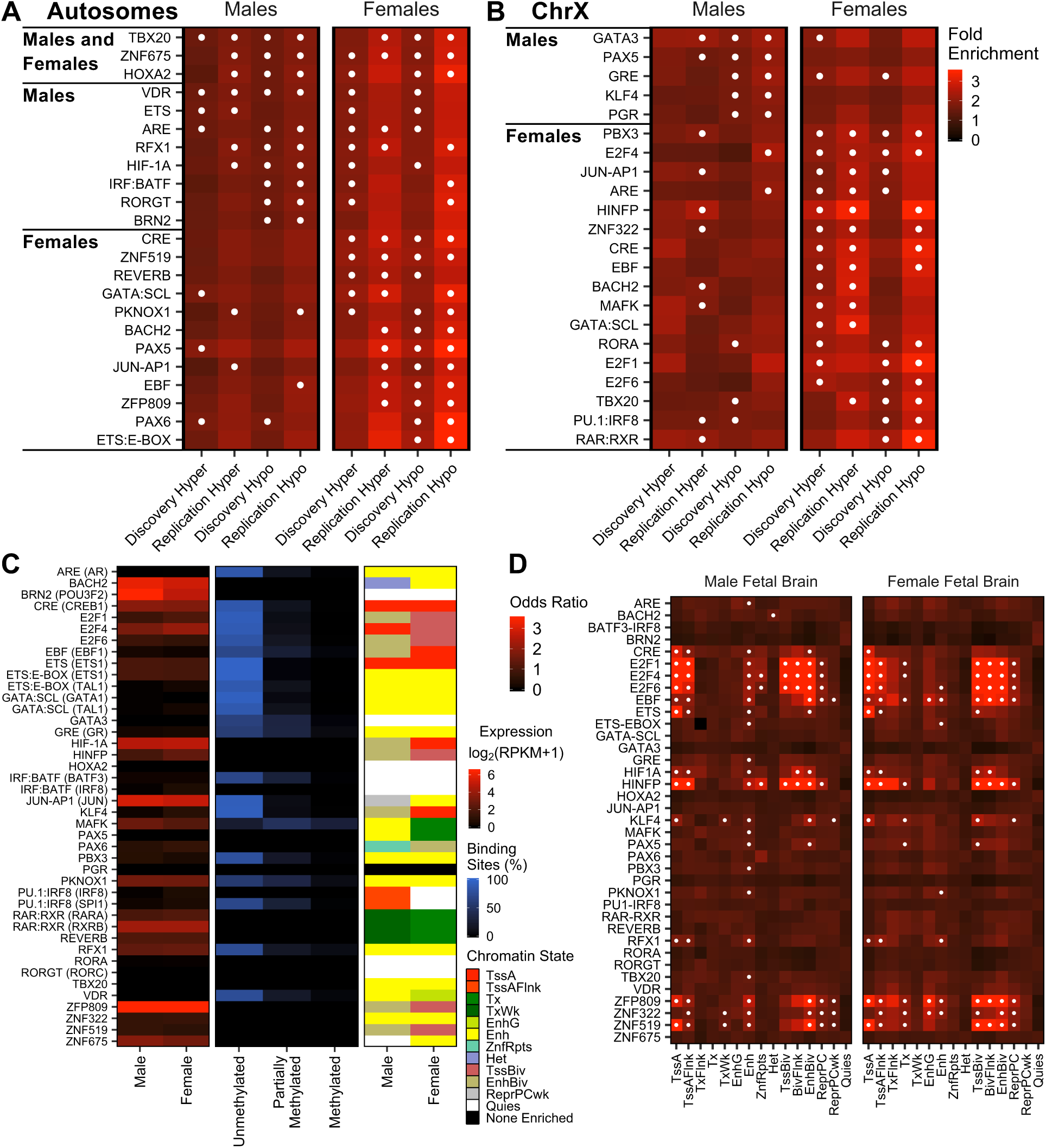
Cord blood ASD DMRs are significantly enriched for motifs for fetal brain-relevant methyl-sensitive transcription factors. Sequences in ASD DMRs were analyzed for enrichment in known transcription factor binding motifs compared to background regions using the Hypergeometric Optimization of Motif EnRichment (HOMER) tool. The fold enrichment for top enriched motifs was plotted for the indicated DMR sets for (A) DMRs on autosomes or (B) DMRs on chromosome X. Motifs were ordered by replicated sex and direction and plotted on the same scale. Top enriched (•) motifs were identified as those with a *q*-value less than 0.05 and present in the top quartile of fold enrichment and -log(*q*-value) within that DMR set. Plotted motifs were the top enriched in both discovery and replication sets of DMRs for that sex and direction. (C) Expression of transcription factors with top enriched motifs in male or female fetal brain; proportion of ChIP-seq peaks in unmethylated (less than 10%), partially methylated (10 - 90%), and methylated (greater than 90%) contexts; and top ranked chromatin state by mean rank of odds ratio and *p*-value for motif in male or female fetal brain and with a *q*-value less than 0.05. (D) Odds ratios for enrichment of top enriched motif locations in male or female fetal brain chromatin states. Top enriched (•) chromatin states were identified as those with a *q*-value less than 0.05 and present in the top quartile of odds ratio and -log(*q*-value) within that tissue (pooled males TD *n* = 56, ASD *n* = 56; pooled females TD *n* = 20, ASD *n* = 20). RPKM, reads per kilobase per million reads;

Among autosomal DMRs in males, hypermethylated DMRs were enriched for ETS and VDR motifs, while hypomethylated DMRs were also enriched for VDR as well as ARE, BRN2, HIF1A, IRF-BATF, RGX1, and RORGT motifs (*q* < 0.05; Fig. 6A). In contrast, hypomethylated X-linked DMRs in males were enriched for GATA3, GRE, KLF4, PAX5, and PGR motifs (*q* < 0.05; Fig. 6B). Unlike in males, three motifs were in common between female autosomal and X-linked DMRs (hyper: CRE and GATA:SCL; hypo: TBX20; Fig. 6A,B, Additional file 3: Table S30). Still, most of the top enriched motifs were unique for female autosomal versus X-linked ASD DMRs. Specifically, motifs for REVERB and ZNF519 were overrepresented among hypermethylated autosome DMRs, while ZNF519, and also BACH2, CRE, EBF, ETS:E-BOX, JUN-AP1, PAX5, PAX6, PKNOX1, and ZFP809 motifs were overrepresented among hypomethylated autosome DMRs (*q* < 0.05; Fig. 6A). For X-linked DMRs in females, hypermethylated DMRs were enriched in ARE, BACH2, E2F4, EBF, HINFP, JUN-AP1, MAFK, PBX3, and ZNF322 motifs, while hypomethylated DMRs were enriched in PBX3 and E2F4, and also E2F1, E2F6, PU.1:IRF8, RAR:RXR, and RORA motifs (*q* < 0.05; Fig. 6B). The enrichment of largely distinct sets of motifs in ASD DMRs from males and females and from autosomes and the X chromosome suggests mostly separate groups of transcription factor motifs are associated with these DMRs.

To aid in the functional interpretation of these binding motifs in the context of methylation alterations in neurodevelopment, we investigated their transcription factor fetal brain expression, methylation sensitivity, and enrichment for fetal brain chromatin states. Transcription factors expressed in fetal brain with motifs enriched in male autosomal ASD DMRs were *BRN2/POU3F2*, *ETS1*, *HIF1A*, *RFX1*, and *ZNF675* (reads per kilobase per million reads (RPKM) > 1), while those enriched in male X-linked DMRs were not expressed in fetal brain (Fig. 6C). Of these expressed transcription factors, ETS1 and RFX1 prefer unmethylated motifs (less than 10% methylation). Transcription factors with female autosome DMR-enriched motifs that were expressed in fetal brain included *BACH2*, *CREB1*, *ETS1*, *JUN*, *PAX6*, *PKNOX1*, *REVERB*, *RFX1*, *ZNF519,* and *ZNF675* (RPKM > 1). Notably, CREB1, ETS1, JUN, PKNOX1, and RFX1 bind selectively to unmethylated motifs. Among transcription factors with female X-linked DMR-enriched motifs, *BACH2*, *CREB1*, *E2F1*, *E2F4*, *E2F6, HINFP*, *JUN*, *MAFK*, *RARA*, *RXRB*, and *ZNF322* were expressed (RPKM > 1). Out of these expressed transcription factors, CREB1, E2F1, E2F4, E2F6, and JUN prefer unmethylated motifs, while MAFK prefers partially methylated motifs (10 - 90% methylation). ASD DMR-associated transcription factor motifs also significantly overlapped with chromatin states in fetal brain involved in transcriptional regulation, including transcription start sites (TssA, TssAFlnk, and TssBiv) and enhancers (Enh, EnhG, and EnhBiv; Fig. 6D, Additional file 3: Table S31). Together, these results support a functional role for differential methylation patterns identified in cord blood to impact transcription factors relevant to the fetal brain and highlight important sex differences.

## Discussion

In this study, we found evidence that a DNA methylation signature of ASD relative to TD outcome exists in cord blood and that specific regions are consistently differentially methylated despite both technical and individual differences between data sets. Replication was stronger at the level of genes and gene ontologies, rather than individual DMRs, although both were significantly higher than expected by chance. For instance, 537 genes in males and 1762 genes in females were consistently associated with DMRs, while 7 DMRs in males and 31 DMRs in females were regionally replicated. Furthermore, replicated cord blood ASD DMR genes identified by our WGBS-based approach across two sequencing platforms significantly overlapped with genes identified from prior epigenetic ASD studies in brain and other tissues using array-based approaches. Together, these results demonstrate a convergence of common dysregulated genes and pathways and suggest that the methylation state of individual CpGs or specific CpG clusters is less replicable than the convergent epigenomic signature. In this way, the cord blood methylome ASD signature likely reflects the consequences of dysregulating functionally related sets of genes in early prenatal development, which is likely more important for the resulting phenotype than precisely altered CpGs.

Implicating their potential etiological relevance, cord blood ASD DMR genes were enriched for expression in brain, at the postsynaptic membrane, and during embryonic development in both males and females. Genes previously identified in epigenetic studies of ASD were also overrepresented among cord blood DMR genes. Of these, *RBFOX1* is particularly interesting, as it has been previously associated with ASD in post-mortem brain by studies of DNA methylation [21, 25], histone acetylation [26], and gene expression [12,38,39], as well as in genome-wide association studies [40]. *RBFOX1* encodes for a neuronal splicing factor, which was identified as the hub gene of an ASD-associated coexpression module in ASD post-mortem brain [41], and whose target genes are enriched among genes contributing to ASD risk [42].

Notably, *RBFOX1* was also found to be differentially methylated in post-mortem brain of subjects with Rett syndrome and Dup15q syndrome, suggesting a fundamental role in neurodevelopment [23]. In contrast, cord blood ASD DMR genes were not enriched for those identified in transcriptome studies of ASD. This, together with the enrichment of cord DMR genes for embryonic expression, suggests the ASD-associated differential methylation identified in cord blood is more likely to be a remnant of past dysregulation in early development rather than a current correlate of transcript levels. The genes most likely to contribute to ASD are predicted to be expressed both in early prenatal development and in brain, which is indeed what we observe in our cord blood ASD DMR genes.

In a complementary approach to gene-level analyses, we investigated the region-based enrichment of ASD DMRs for CpG context, histone PTMs, chromatin states, and transcription factor motifs, and found similar epigenomic signatures for autosome DMRs in males and females. Overall, *cis*-regulatory regions present in almost all tissues were overrepresented among autosome DMRs, including transcription start sites, bivalent regions, and polycomb-repressed regions. Many transcription factors associated with ASD DMRs were specific to males or females, sensitive to methylation, and expressed in fetal brain. Specifically, ETS1, RFX1, and CREB1 were enriched in specific fetal brain chromatin states, and all three displayed a sexually-dimorphic pattern of enrichment in ASD DMRs, suggesting they may be involved in sex-specific transcriptional dysregulation in ASD. ETS1 is widely expressed during embryogenesis and is involved in organ formation and morphogenesis of the mesoderm [43]. RFX1 is necessary for the formation of the testis cord in the fetus and plays a role in spermatogenesis [44]. CREB1 is important for neuronal stimulus-dependent gene expression, and its deletion in mice results in impaired survival of sensory and sympathetic neurons and axonal elongation [45]. Together, the functional implications of these findings are that cord blood ASD DMRs reflect early perturbations in neurodevelopment. Although ASD DMRs on autosomes are associated with similar chromatin states in both sexes, affected transcription factor binding sites are sex-specific.

Cord blood ASD DMRs identified from both sexes were enriched for X-linked genes and for distinctive chromatin states on the X chromosome compared to autosomes. Replicated ASD DMR genes on the X chromosome included 21 genes common to males and females, 34 only in males, and 152 only in females, with most of these genes previously associated with ASD in at least one genome-wide study. Compared to all genes in the genome, genes on the X chromosome are enriched for expression in brain and embryonic development and also mental retardation, suggesting that chromosome-wide epigenetic dysregulation of the X chromosome may predispose individuals to neurodevelopmental disorders. Additional evidence for this premise can be seen in X chromosome aneuploidy disorders such as Turner, Klinefelter, and trisomy X syndromes, where differences in neuroanatomy and behavior have been identified [46–51]. Interestingly, both Klinefelter and trisomy X subjects are at a more than four-fold increased risk of having ASD [51]. In addition, a large number of X-linked disorders involve intellectual disability [52].

X-linked ASD DMRs exhibited distinct chromatin and transcription factor features from those on autosomes, with notable sex differences. Male X-linked ASD DMRs were less enriched for active regulatory elements, but more enriched for quiescent regions compared to autosomal ASD DMRs. In contrast, female X-linked ASD DMRs were more enriched in transcriptionally-active regions than autosomal ASD DMRs. Similar patterns were observed regardless of the direction of methylation change or tissue type. Additionally, only female X-linked ASD DMRs were enriched for the E2F1, E2F4, and E2F6 transcription factors, which are all expressed in fetal brain, sensitive to methylation, and regulate cell cycle progression [53]. The disruption of active regulatory regions in females can also be seen in the large number of replicated X chromosome DMR genes in females compared to males, including *MECP2*, which was previously found to have altered methylation in female ASD brain [54]. Together, these results demonstrate that female-specific X-linked ASD DMRs predominate in euchromatic regions, while those specific to males are predominantly heterochromatic, suggesting sex differences due to epigenetic mechanisms on the X chromosome.

One potential explanation for this distinctive female pattern of epigenetic dysregulation could be the phenomenon of “X chromosome erosion”, recently identified in female pluripotent stem cells [55]. During the process of random X chromosome inactivation that occurs in human peri-implantation embryos, the long noncoding RNA *XIST* coats all X chromosomes in males and females, while another primate-specific X-linked long noncoding RNA named *XACT* coats only active X chromosomes to prevent transcriptional repression [56]. When X chromosome inactivation becomes eroded in cultured human embryonic stem cells, *XACT* is aberrantly expressed, resulting in decreased *XIST*, loss of repressive H3K27me3 and DNA methylation at promoters, and transcriptional reactivation of some genes on the inactive X [55]. Notably, *XACT* was associated with ASD DMRs in both males and females, and its expression is normally exclusive to a preimplantation developmental window between 4-8 cells and the epiblast. We therefore hypothesize that aberrant *XACT* expression and/or X chromosome erosion during early development could underlie our findings of X-linked epigenetic dysregulation in cord blood from newborns later diagnosed with ASD. A second X chromosome in females may then serve as a mechanism of epigenetic protection against ASD and contribute to the observed 3 to 1 male bias in ASD [14].

A balanced interpretation of the results of this study should be guided by its strengths and limitations. The strengths of this study lie in its design and approach. Subjects were obtained from two prospective high-familial risk cohorts that were deeply phenotyped during the first three years of life, which included the gold-standard ADOS assessment. The high-familial risk design increases ASD prevalence in a prospective cohort, as siblings of an ASD proband are significantly more likely to develop ASD or other developmental delays [57, 58]. Notably, this type of design also increases the power to detect environmental factors contributing to ASD risk [59]. In this instance, it is possible that the high-familial risk design could increase the ability to capture changes in DNA methylation related to either inherited early developmental programming or a marker of environmentally-induced risk. In our analytical approach, we used WGBS to assay DNA methylation at more than 20 million CpGs, and stratified subjects for DMR identification by both sex and sequencing platform, which were two major drivers of variability in methylation. Furthermore, we focused our analysis on systems-level features informed by chromosome biology, including gene locations and functions, as well as chromatin states and transcription factors at *cis*-regulatory regions. Findings were confirmed by replication in two groups of participants, different sequencing platforms, different bioinformatic approaches, and prior methylation studies of ASD in other tissues and platforms.

The limitations of this study include the heterogeneous nature of both umbilical cord blood DNA and ASD etiology, as well as the inherent limitations of sample size and dichotomous ASD outcome in this prospective study design. Additionally, findings in these high-familial risk cohorts may not translate to low-risk populations. Cord blood is a complex tissue with dozens of different cell types that are undergoing dynamic changes in the prenatal period. Because we analyzed this tissue in bulk, the DMRs identified could reflect methylation alterations within individual cell types as well as changes in the proportions of these cells between subjects, potentially reducing power to detect disease relevant signatures. Notably, we identified an increased proportion of nRBCs in males with ASD, which correlated with lower global CpG methylation; however, nRBCs only correlated with methylation at a small number of ASD DMRs. Our demonstration of a lack of strong correlation of methylation within ASD DMRs with estimated cell proportions combined with the replication in ASD brain studies and tissue-independence of the *cis*-regulatory regions identified alleviate these important concerns. Additionally, ASD is a heterogeneous disorder with high variability between subjects in genetic and environmental risk factors and comorbid phenotypes, which together with the small sample size and dichotomous outcome, may have limited our ability to detect small methylation differences in individual genes or CpG sites at high confidence; however, for those identified differences, we verified associations between methylation and continuous ASD symptom severity and cognitive functioning with more power, and confirmed the lack of associations with non-neurodevelopmental factors. We supplemented this approach by identifying regions with empirical *p*-value significance across two groups of subjects to reduce false positives and focused on convergent genes, gene pathways, and chromatin-level features.

### Conclusions

In this first study of the entire cord blood methylome from newborns later diagnosed with ASD, we identified differential methylation at birth at multiple levels, including regions, genes, functional gene sets, and chromatin states, and replicated these findings across two groups. In both males and females, ASD DMR associated genes were more likely to be located on the X chromosome, to be expressed in brain and at the postsynaptic membrane, and to modify transcription factor binding sites relevant to the developing brain. Autosomal ASD DMRs were also enriched for *cis*-regulatory functional regions present across most tissues, including transcription start states, CpG islands, enhancers, and bivalent regions. ASD DMRs on the X chromosome reflected chromosome- and sex-specific dysregulation, with an enrichment for quiescent regions among male DMRs, and an enrichment in open chromatin states in female DMRs, compared to those on autosomes. These findings in cord blood suggest that epigenetic dysregulation in ASD may originate during early prenatal development in a sex-specific manner and converge on brain-relevant genes to disrupt neurodevelopment.

## Methods

### Sample population and biosample collection

#### MARBLES

The MARBLES study recruited mothers of children receiving services through the California Department of Developmental Services for their diagnosis of ASD; mothers must have been either planning a pregnancy or already pregnant with another child. Study inclusion criteria were: the mother or father has at least one biological child with ASD; the mother is at least 18 years old; the mother is pregnant; the mother speaks, reads, and understands English at a sufficient level to complete the protocol, the younger sibling will be taught to speak English; and the mother lives within 2.5 hours of the Davis/Sacramento region at the time of enrollment. As previously described in depth [60], demographic, lifestyle, environmental, diet, and medical information were collected prospectively through telephone-assisted interviews and mailed questionnaires during pregnancy and the postnatal period. Mothers were provided with umbilical cord blood collection kits prior to delivery. Arrangements were made with obstetricians/midwives and birth hospital labor/delivery staff to ensure proper sample collection and temporary storage. As described below, infants received standardized neurodevelopmental assessments between six months and three years of age.

#### EARLI

The EARLI study recruited and followed pregnant mothers who had an older child diagnosed with ASD from pregnancy through the first three years of life and has been described in detail previously [61]. EARLI families were recruited at four EARLI network sites (Drexel/Children’s Hospital of Philadelphia, Johns Hopkins/Kennedy Krieger Institute, Kaiser Permanente Northern California, and University of California, Davis) in three US regions (Southeast Pennsylvania, Northeast Maryland, and Northern California). In addition to having a biological child with ASD confirmed by EARLI study clinicians, inclusion criteria also consisted of being able to communicate in English or Spanish; being 18 years or older; living within two hours of a study site; and being less than 29 weeks pregnant. EARLI research staff made arrangements with obstetricians/midwives and birth hospital labor/delivery staff to ensure proper umbilical cord blood sample collection and temporary storage. The development of children born into the cohort was closely followed using standardized neurodevelopmental assessments through three years of age.

### Diagnostic classification

In both the MARBLES and EARLI studies, child development was assessed by trained, reliable examiners. Diagnostic assessments at 36 months included the gold standard ADOS [62], the Autism Diagnostic Interview-Revised [63], conducted with parents, and the MSEL [64], a test of cognitive, language, and motor development. Based on a previously published algorithm that uses ADOS and MSEL scores [65, 66], children included in the study were classified into one of three exclusive outcome groups: ASD, TD, or Non-typically developing; however, the analyses for this paper were limited to samples from children with ASD and TD outcomes. Children with ASD outcomes had scores over the ADOS cutoff and met DSM-5 criteria for ASD. Children with TD outcomes had all MSEL scores within 2 SD of the average and no more than one MSEL subscale 1.5 SD below the normative mean and scores on the ADOS at least three points below the ASD cutoff.

### Demographic characteristics

In both studies, demographic information was collected prospectively throughout gestation and the postnatal period with in-person and telephone-assisted interviews and mailed questionnaires. Demographic characteristics were tested for differences relative to diagnostic outcome across all subjects using Fisher’s exact test for categorical variables and one-way ANOVA for continuous variables. *P*-values were adjusted for the total number of demographic variables using the false discovery rate (FDR) method [67].

### WGBS sample processing

DNA was extracted from whole umbilical cord blood with the Gentra Puregene Blood kit (Qiagen, Hilden, Germany) and quantified with the Qubit dsDNA HS Assay kit (Thermo Fisher Scientific, Waltham, MA, USA). DNA was bisulfite converted with the EZ DNA Methylation Lightning kit (Zymo, Irvine, CA, USA). Sodium bisulfite treatment converts all unmethylated cytosine to uracil residues, which are detected as thymine after library preparation [68]. WGBS libraries were prepared from 100ng of bisulfite-converted DNA using the TruSeq DNA Methylation kit (Illumina, San Diego, CA, USA) with indexed PCR primers and a 14-cycle PCR program. Libraries for the discovery sample set were sequenced at 2 per lane with 150 base pair paired-end reads and spiked-in PhiX DNA on the HiSeq X Ten (Illumina, San Diego, CA, USA) by Novogene (Sacramento, CA, USA). Samples included in the discovery set were obtained from 74 males (TD *n* = 39, ASD *n* = 35) and 32 females (TD *n* = 17, ASD *n* = 15) in the MARBLES and EARLI studies (MARBLES *n* = 42, EARLI *n* = 64). Libraries for the replication sample set were sequenced over two lanes indexed at 4 samples per lane with 100 base pair single-end reads and spiked-in PhiX DNA on the HiSeq 4000 (Illumina, San Diego, CA, USA) by the Vincent J. Coates Genomics Sequencing Laboratory at University of California, Berkeley. Samples included in the replication set were obtained from 38 Males (TD *n* = 17, ASD *n* = 21) and 8 Females (TD *n* = 3, ASD *n* = 5) in the MARBLES study.

### WGBS read alignment and quality control

Sequencing reads were preprocessed, aligned to the human genome, and converted to CpG methylation count matrices with CpG_Me (v1.0, [69–71]) using the appropriate pipeline for single- or paired-end reads. After Illumina quality filtering of raw fastq files, reads were trimmed to remove adapters and both 5’ and 3’ methylation bias. Trimmed reads were screened for contaminating genomes, aligned to the hg38 human reference genome assembly, and filtered for PCR duplicates. Counts of CpG methylation at all covered sites were extracted to generate Bismark cytosine methylation reports. Comprehensive quality control reports were examined for each sample, and libraries with incomplete bisulfite conversion were excluded, as measured by CHH methylation greater than 2%. Sex was confirmed for each sample based on coverage of the X and Y chromosomes. The CpG_Me workflow incorporates the Trim Galore! (v0.4.5, RRID:SCR_011847, [72]), Bismark (v0.19.1, RRID:SCR_005604, [70]), Bowtie 2 (v2.3.4.1, RRID:SCR_005476, [73]), FastQ Screen (v0.11.4, RRID:SCR_000141, [74]), SAMtools (v1.8, RRID:SCR_002105, [75]), and MultiQC (v1.5, RRID:SCR_014982, [76]) packages.

### Global DNA methylation analysis and covariate associations

Global CpG methylation for each sample was extracted from Bismark count matrices as the total number of methylated CpG counts divided by the total CpG counts across all chromosomes. Global methylation was compared with behavioral, demographic, and technical variables using linear regression, stratified by sex, and examined both within and pooled across sequencing platforms. Linear models included adjustment for PCR duplicates and also sequencing platform when combined. Continuous variables were converted to SD before linear regression. *P*-values were adjusted for the number of variables using the FDR method.

### Cell-type deconvolution

Cell-type proportions were deconvoluted from WGBS methylation data based on a umbilical cord blood reference panel, which defined cell-type-specific CpG sites in B cells, CD4+ and CD8+ T cells, granulocytes, monocytes, natural killer cells, and nucleated red blood cells using the Infinium HumanMethylation450 BeadChip array [35]. Percent methylation was extracted from WGBS Bismark methylation count matrices at cell-type-specific loci covered in at least 90% of samples, which included between 76 and 85 CpG sites per cell type. Missing values were imputed with the mean percent methylation for that locus across all samples. Percent methylation data, along with model parameters defined in the reference panel, were input into the projectCellType() function in the minfi R package (v1.28.4, RRID:SCR_012830, [77]) to estimate cell-type proportions, which were scaled to 100% for each sample.

### DMR and DMB identification

DMRs were identified between ASD and TD subjects in only males, only females, and all subjects with adjustment for sex. DMBs were identified between ASD and TD subjects within sex only. The same comparisons for DMRs and DMBs were done in the discovery and replication sets. DMRs and DMBs were identified using the DMRichR (v1.0, [69, 78]), dmrseq (v1.2.3, [79]), and bsseq (v1.18.0, RRID:SCR_001072, [36]) R packages. The beta-binomial distribution, variance in percent methylation, and spatial correlations inherent in WGBS data can be appropriately modeled using a generalized least squares regression model with a nested autoregressive correlated error structure as implemented in the dmrseq package [79, 80]. In this approach, candidate regions are identified based on consistent differences in mean methylation between groups, and region-level statistics are estimated which account for coverage, mean methylation, and correlation between CpGs. These statistics are then compared to a permutation-generated pooled null distribution to calculate an empirical *p*-value. In this study, Bismark count matrices were filtered for CpGs covered by at least 1 read in 50% of samples in both groups, but if less than 10 samples were present in a group, the threshold was increased to 1 read in all samples. DMRs were identified with the dmrseq() function from the dmrseq package using the default parameters, except the single CpG methylation difference coefficient cutoff was set to 0.05 and the minimum number of CpGs was set to 3. DMRs were called if the permutation-based empirical *p*-value was less than 0.05. DMBs were also identified with the dmrseq() function using the default parameters for blocks, except the single CpG methylation difference coefficient cutoff was set to 0.01 and the minimum number of CpGs was set to 3. As described in the dmrseq package vignette [79], the default parameters for DMBs differ from those for DMRs in that the minimum width for DMBs is 5 kilobases and the maximum gap between CpGs in a DMB is also 5 kilobases. Additionally, the smoothing span window is widened by setting the minimum CpGs in a smoothing window to 500, the width of the window to 50 kilobases, and the maximum gap between CpGs in the same smoothing cluster to 1 megabase. Background regions were defined as the genomic locations where it is possible to identify a DMR, which were those regions in the coverage-filtered Bismark count matrix with at least 3 CpG sites less than 1 kilobase apart.

### DMR hierarchical clustering and principal component analysis (PCA)

The ability of DMRs to distinguish between ASD and TD subjects was tested through clustering subjects based on percent methylation in DMRs using hierarchical clustering and PCA. In both approaches, percent methylation in DMRs was extracted from Bismark methylation count matrices as the number of methylated CpG counts divided by the total CpG counts in that DMR. For hierarchical clustering, the mean percent methylation in a DMR across all subjects was subtracted from the percent methylation in a DMR for each subject. Both subjects and DMRs were clustered using the Euclidean distance and Ward’s agglomeration method. For PCA, missing values were imputed with the mean percent methylation in that DMR. PCA was performed using the prcomp() function in the stats R package with centering to zero and scaling to unit variance, and then visualized using the ggbiplot R package [81] with the ellipse indicating the 95% confidence interval for each group.

### DMR-covariate associations

Percent methylation in DMRs was extracted from Bismark methylation count matrices as the number of methylated CpG counts divided by the total CpG counts in that DMR. Continuous variables were converted to SD, and methylation at each DMR was compared with behavioral, demographic, and technical variables using linear regression. *P*-values were adjusted for the number of variables and DMRs using the FDR method.

### Computational validation of DMRs

DMRs from all comparisons conducted within sex were examined for validation by recalling them using the BSmooth.tstat() and dmrFinder() functions from the bsseq R package (v1.18.0, RRID:SCR_001072, [36]), which uses a different statistical approach. In this method, percent methylation values for individual CpGs are smoothed to incorporate information from neighboring loci. T-tests are conducted at each CpG site to compare the two groups, and adjacent CpGs with *p*-values less than 0.05 are grouped into DMRs. Default parameters were used for BSmooth.tstat(), except the variance was estimated assuming it was the same for both groups. Default parameters were also used for dmrFinder(), except the t-statistic cutoff was set to the value where *p* = 0.05 in a two-sided t-test with *n* - 2 degrees of freedom, according to the qt() function. DMRs were further filtered for at least 3 CpGs, average methylation difference greater than 0.05, and inverse density of at most 300. The genomic locations of the DMRs from the two different methods and with the same direction of methylation difference were overlapped. The significance of the overlap between the DMR sets was tested using a permutation-based test implemented with the regioneR R package (v1.14.0, [82]). Both sets of DMRs were redefined as the set of background regions containing a DMR for that set. Empirical *p*-values were calculated by comparing the true overlap to a null distribution of overlaps between 10,000 length-matched region sets randomly sampled from the background.

### DMR replication by region overlap, differential methylation, and gene overlap

DMRs identified within sex between ASD and TD subjects in the discovery sample set were examined for replication in the independent replication sample set using three different methods: region overlap, differential methylation, and gene overlap. In the region overlap method, DMRs were identified separately for samples in the discovery and replication sets and the genomic locations of DMRs with the same direction of methylation difference were overlapped. Significance of the overlap between the discovery and replication DMRs was tested using a permutation-based test implemented with the regioneR R package (v1.14.0, [82]) as with the computational validation of the DMRs, except the background regions from the discovery and replication sets were intersected to generate a consensus background.

In the differential methylation replication method, percent methylation at DMRs identified in discovery samples was first extracted from Bismark count matrices for both discovery and replication samples. DMRs not covered in more than 50% of either group in the replication sample set were excluded. Percent methylation at each DMR was compared with diagnosis outcome separately in discovery and replication samples using linear regression, and replicated DMRs were called as those with the same direction and an unadjusted *p*-value less than 0.05 in both sample sets.

In the gene overlap method, genes were first annotated to DMRs using the same approach employed by GREAT [83], but adapted for the hg38 assembly. In this strategy, every gene in the NCBI RefSeq database is given a regulatory domain, where the basal regulatory domain extends 5 kilobases upstream and 1 kilobase downstream of the transcription start site (TSS), regardless of other genes. The basal regulatory domain is then extended upstream and downstream of the TSS to the nearest gene’s proximal domain or 1 megabase, whichever is closer. Finally, a gene is assigned to a DMR if the DMR overlaps its regulatory domain. All genes annotated to discovery DMRs for that sex were overlapped with all genes annotated to replication DMRs for that sex. The significance of DMR gene overlap between the discovery and replication sets was tested using Fisher’s exact test implemented with the GeneOverlap R package (v1.18.0, [84], and compared to genes annotated to both sets of background regions. Replicated DMR genes were defined as those annotated to both discovery and replication DMRs for that sex. Replicated DMR genes in males were tested for significant overlap with replicated DMR genes in females as above. DMB genes were also examined for replication with the gene overlap method.

### Gene functional enrichment

ASD DMR gene functional enrichment was assessed using the Database for Annotation, Visualization, and Integrated Discovery (DAVID, v6.8, RRID:SCR_001881, [85]) and the accompanying RDAVIDWebService R package (v1.20.0, [86]). For both discovery and replication sets, enrichments of genes annotated to sex-stratified ASD DMRs were compared with genes annotated to background regions, and these were input into DAVID as NCBI Entrez ID numbers. *P*-values were adjusted using the Benjamini method and significantly enriched terms were called as those with *q*-values less than 0.05. Enriched terms were defined as replicated if they were significantly enriched in both discovery and replication DMR genes. Functional enrichment of DMB genes was assessed as above. Functional enrichments of all X chromosome genes were also done similarly, with X-linked RefSeq genes compared to all RefSeq genes using their Entrez IDs.

### Autism gene enrichment

For both discovery and replication sets, enrichment of sex-stratified ASD DMR genes with those identified in previous genomic, epigenomic, and transcriptional studies of ASD was examined. Gene lists were sourced from the reported hits of each paper [6,12,21–30,37–40,87–103]. Reported regions from epigenomic studies were converted to the hg38 assembly using the liftOver() function from the rtracklayer R package (v1.42.2, [104]), and genes were annotated using the same approach used in GREAT [83]. After converting to Entrez IDs, overlap significance was tested with Fisher’s exact test using the GeneOverlap R package (v1.18.0, [84]) and compared to genes annotated to background regions. P-values were adjusted for the number of gene lists using the FDR method, and significant overlaps were called as those with *q*-values less than 0.05. Replicated overlaps were identified as the gene lists that significantly overlapped with both discovery and replication DMR genes.

### CpG context, ChIP-seq, and ChromHMM chromatin state enrichments

For both discovery and replication sets, sex-stratified ASD DMRs were assessed for region-based enrichment with CpG context, histone PTM ChIP-seq peaks, and ChromHMM-defined chromatin states using the Locus Overlap Analysis (LOLA) R package (v1.12.0, [105]). CpG context maps were obtained from the annotatr R package (v1.8.0, [106]), while histone ChIP-seq peaks and chromatin states were obtained from the Roadmap Epigenomics Project (RRID:SCR_008924, [107]). As recommended in the LOLA documentation [105], enrichments were conducted with DMRs redefined as the set of background regions containing a DMR compared to the set of all background regions. For appropriate visualization in heatmaps, overlaps with less than 5 regions were excluded, and infinite odds ratios and *p*-values were replaced with the maximum value for that DMR set. *P*-values were adjusted with the FDR method, and top enriched regulatory regions were defined as those where the *q*-value was less than 0.05 and both the odds ratio and -log_10_(*q*-value) were at least the median value for more than half of the 127 cell types examined. Significance of differential enrichment of X chromosome compared to autosome ASD DMRs was assessed by paired two-sided t-test of odds ratios for each cell type for that regulatory region. *P*-values were adjusted for the number of different regulatory regions with the FDR method and significant differential enrichment was called if the *q*-value was less than 0.05.

### Transcription factor motif enrichment

For both discovery and replication sets, enrichment for known transcription factor binding site motif sequences in sex-stratified ASD DMRs was examined using the Hypergeometric Optimization of Motif EnRichment (HOMER) tool (v4.10, RRID:SCR_010881, [108]). After redefining DMRs as the subset of background regions containing a DMR, DMRs and background regions were input into findMotifsGenome.pl with the default parameters, except the region size was set to given, sequences were normalized for percent CpG content, and the number of randomly sampled background regions was increased to 100,000. Fold enrichment was calculated as the percent of DMR sequences with the motif divided by the percent of background sequences with the motif. *P*-values were adjusted for the number of known motifs tested using the FDR method. Top enriched motifs were defined as those with a *q*-value less than 0.05 and both fold enrichment and -log_10_(*q*-value) in at least the top quartile for that DMR set. Replicated motifs were those identified as top enriched in both discovery and replication DMR sets for that direction.

### Replicated transcription factor motif analysis

Transcription factors that replicated as among the top enriched in sex-stratified ASD DMRs across the discovery and replication sample sets were investigated for motif enrichment in fetal brain chromatin states, expression in fetal brain, and methylation sensitivity. To test for motif enrichment in chromatin states in fetal brain, genomic locations for replicated motif sequences were first determined with scanMotifGenomeWide.pl from HOMER (v4.10, RRID:SCR_010881, [108]). Background regions were defined as the set of regions in the genome where any of the known motifs tested in HOMER were present. Enrichment in male or female fetal brain chromatin states from the Roadmap Epigenomics Project (RRID:SCR_008924, [107]) was tested using the LOLA R package (v1.12.0, [105]), with replicated motif locations redefined as a subset of the background regions containing a motif. P-values were adjusted using the FDR method, and top enriched chromatin states were those with a *q*-value less than 0.05, and with both odds ratio and -log_10_(*q*-value) in at least the top quartile for that tissue type. The top ranked chromatin state for each motif was the top enriched state with the lowest mean rank of odds ratio and *p*-value. RNA-seq expression data of replicated transcription factors in fetal brain were obtained from the Allen BrainSpan Atlas of the Developing Human Brain (RRID:SCR_008083, [109]). Displayed data were derived from the dorsolateral prefrontal cortex from one male (ID# 12820) and one female (ID# 12834) donor at 13 weeks post conception. RNA-seq expression data of replicated X-linked DMR genes were also derived from this source. Methylation sensitivity data were obtained from MethMotif (v1.3, [110]), and counts for each motif corresponding to unmethylated (less than 10%), partially methylated (10 - 90%), and methylated (greater than 90%) loci were summed and divided by the total counts for that motif to determine the proportion of binding sites in each methylation category.

## Abbreviations

Ac, acetylation; ADOS, Autism Diagnostic Observation Schedule; ASD, autism spectrum disorder; BivFlnk, flanking bivalent transcription start site or enhancer; ChIP-seq, chromatin immunoprecipitation sequencing; chr, chromosome; DAVID, Database for Annotation, Visualization, and Integrated Discovery; DEG, differentially-expressed genes; DMB, differentially-methylated block; DMR, differentially-methylated region; Dup15, Chromosome 15q11-q13 Duplication syndrome; EARLI, Early Autism Risk Longitudinal Investigation; Enh, enhancer; EnhBiv, bivalent enhancer; EnhG, genic enhancer; EWAS, epigenome-wide association study; FDR, false discovery rate; H3K27, histone 3 lysine 27; H3K36, histone 3 lysine 36; H3K4, histone 3 lysine 4; H3K9, histone 3 lysine 9; Het, heterochromatin; HOMER, Hypergeometric Optimization of Motif EnRichment; hyper, hypermethylated; hypo, hypomethylated; LCL, lymphoblastoid cell line; LOLA, Locus Overlap Analysis; MARBLES, Markers of Autism Risk in Babies - Learning Early Signs; mCpG, CpG methylation; mCpH, CpH methylation; me3, trimethylation; MSEL, Mullen Scales of Early Learning; nRBC, nucleated red blood cell; PFC, prefrontal cortex; PTM, post-translational modification; Quies, quiescent region; ReprPC, polycomb-repressed region; ReprPCWk, weak polycomb-repressed region; RPKM, reads per kilobase per million reads; RTT, Rett syndrome; SD, standard deviation; SFARI, Simons Foundation Autism Research Initiative; TD, typically developing; TSS, transcription start site; TssA, active transcription start site; TssAFlnk, flanking active transcription start site; TssBiv, bivalent transcription start site; Tx, strong transcription; TxFlnk, transcribed at gene 5’ and 3’; TxWk, weak transcription; WGBS, whole-genome bisulfite sequencing; ZnfRpts, zinc finger genes and repeats;

## Declarations

### Ethics approval and consent to participate

The University of California, Davis Institutional Review Board and the State of California Committee for the Protection of Human Subjects approved this study and the MARBLES Study protocols. Human Subjects Institutional Review Boards at each of the four sites in the EARLI Study approved this study and the EARLI Study protocols. Neither data nor specimens were collected until written informed consent was obtained from the parents.

### Consent for publication

Not applicable.

### Availability of data and material

The datasets supporting the conclusions of this article are available in the Gene Expression Omnibus repository at accession number GSE140730 (https://www.ncbi.nlm.nih.gov/geo/query/acc.cgi?acc=GSE140730). Data are shared if parents gave informed consent. All code used in the analyses for this study is available on GitHub (https://github.com/cemordaunt/AutismCordBloodMethylation).

### Authors’ contributions

CEM was the lead author and contributed substantially to DNA extraction, library preparation, all bioinformatic data analysis, data visualization, interpretation of results, and writing of the manuscript. JMJ provided substantial assistance with sample organization, DNA extraction, library preparation, and bioinformatic data analysis. BL contributed extensively to sequence alignment, quality control, and WGBS data analysis. YZ, KWD, and KMB added critical advice for bioinformatic data analysis methods and interpretation. JIF contributed to sample handling and organization. HEV, KL, LAC, CJN, SO, IH, MDF, and RJS contributed to study design and also subject acquisition, diagnosis, and characterization. JML and RJS conceived of the study and funded the whole-genome bisulfite sequencing. JML contributed substantially to data interpretation and revision of the manuscript. All authors read and approved the final manuscript.

## Acknowledgements

We would like to sincerely thank the participants in the MARBLES and EARLI studies.

