## Additional File 1 - Supplemental Figures for "Cord blood DNA methylome in newborns later diagnosed with autism spectrum disorder reflects early dysregulation of neurodevelopmental and X-linked genes"

### Table of Contents


**Lower global DNA methylation in umbilical cord blood from males later diagnosed with ASD corresponds with increased nucleated red blood cells** **3**

Supplemental Figure 1. Global CpG methylation is associated with diagnosis and behavioral outcome scores only in males 3

Supplemental Figure 2. The proportion of nRBCs is associated with behavioral outcome and global CpG methylation in males 4

**Region-specific differential methylation patterns in umbilical cord blood distinguish males and females later diagnosed with ASD from typically developing controls** **6**

Supplemental Figure 3. Methylation at ASD DMRs in the discovery set is specifically associated with behavioral outcome 6

Supplemental Figure 4. DMRs identified in all discovery subjects distinguish ASD from TD subjects 7

Supplemental Figure 5. DMRs identified in males and females distinguish ASD from TD subjects in the replication set 9

Supplemental Figure 6. Methylation at ASD DMRs in the replication set is specifically associated with behavioral outcome 10

Supplemental Figure 7. DMRs identified in all replication subjects distinguish ASD from TD subjects 11

Supplemental Figure 8. The majority of ASD DMRs do not overlap probes on the 450K and EPIC arrays 12

**ASD DMRs in umbilical cord blood replicate across independent groups of subjects** **14**

Supplemental Figure 9. A subset of ASD diagnosis DMRs are associated with ASD severity in independent sample sets 14

Supplemental Figure 10. ASD DMRs identified with adjustment for sex miss many genes found when stratifying for sex 15

**Genes in ASD differentially-methylated blocks replicate between independent groups of subjects and are enriched for ASD DMR genes, cadherins, and developmental genes** **16**

Supplemental Figure 11. Replicated ASD DMB genes overlap with replicated ASD DMR genes 16

Supplemental Figure 12. ASD DMB genes are enriched for membrane, cell adhesion, and embryo-expressed genes 17

**Cord blood ASD DMR genes are enriched for neurodevelopmental genes on the X chromosome that are epigenetically dysregulated in ASD brain** **18**

Supplemental Figure 13. Neurodevelopmental genes are overrepresented on the X chromosome 18

Supplemental Figure 14. Replicated DMR genes on the X chromosome are expressed in fetal brain 19

Supplemental Figure 15. Female-specific replicated DMR genes on the X chromosome are expressed in fetal brain 20

Supplemental Figure 16. Cord blood ASD DMR genes are significantly enriched for epigenetically dysregulated genes in ASD brain 22

Supplemental Figure 17. Selected regions with replicated sex-independent DMR genes on the X chromosome 24
Supplemental Figure 18. Selected regions with replicated male-specific DMR genes on the X chromosome 25
Supplemental Figure 19. Selected regions with replicated female-specific DMR genes on the X chromosome 26
**ASD DMRs are enriched for a pan-tissue epigenomic signature that differs between males and females on the X chromosome 27**
Supplemental Figure 20. ASD DMRs in replication subjects are differentially enriched for chromatin states on the X chromosome 27
Supplemental Figure 21. ASD DMRs in discovery subjects are differentially enriched for histone PTMs on the X chromosome 29
Supplemental Figure 22. ASD DMRs in replication subjects are differentially enriched for histone PTMs on the X chromosome 30
Supplemental Figure 23. ASD DMRs on the X chromosome are enriched near CpG islands only in females 31

**Lower global DNA methylation in umbilical cord blood from males later diagnosed with ASD corresponds with increased nucleated red blood cells**

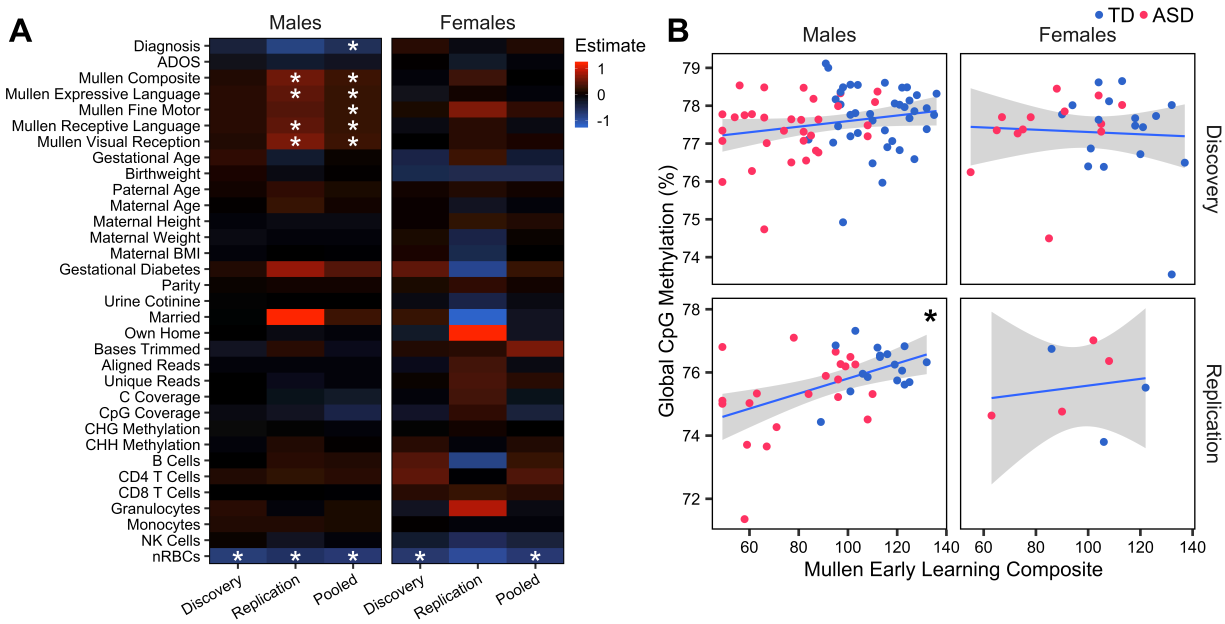


**Supplemental Figure 1. Global CpG methylation is associated with diagnosis and behavioral outcome scores only in males.** (A) Estimated change in global methylation with behavioral, demographic, and technical variables. *P*-values were adjusted for the number of variables using the false discovery rate (FDR) method (* $q < 0.05$). (B) Global CpG methylation compared to Mullen Early Learning Composite score is plotted by sex and sample set (* $p < 0.05$, pooled males $p = 7.9E-4$, pooled females $p = 0.96$). (A,B) Significance was tested using linear regression with adjustment for PCR duplicates, and also adjusted for sequencing platform when pooled (pooled males typically developing (TD) $n = 56$, autism spectrum disorder (ASD) $n = 56$; pooled females TD $n = 20$, ASD $n = 20$). ADOS, Autism Diagnostic Observation Schedule; BMI, body mass index; NK, natural killer; nRBC, nucleated red blood cell;

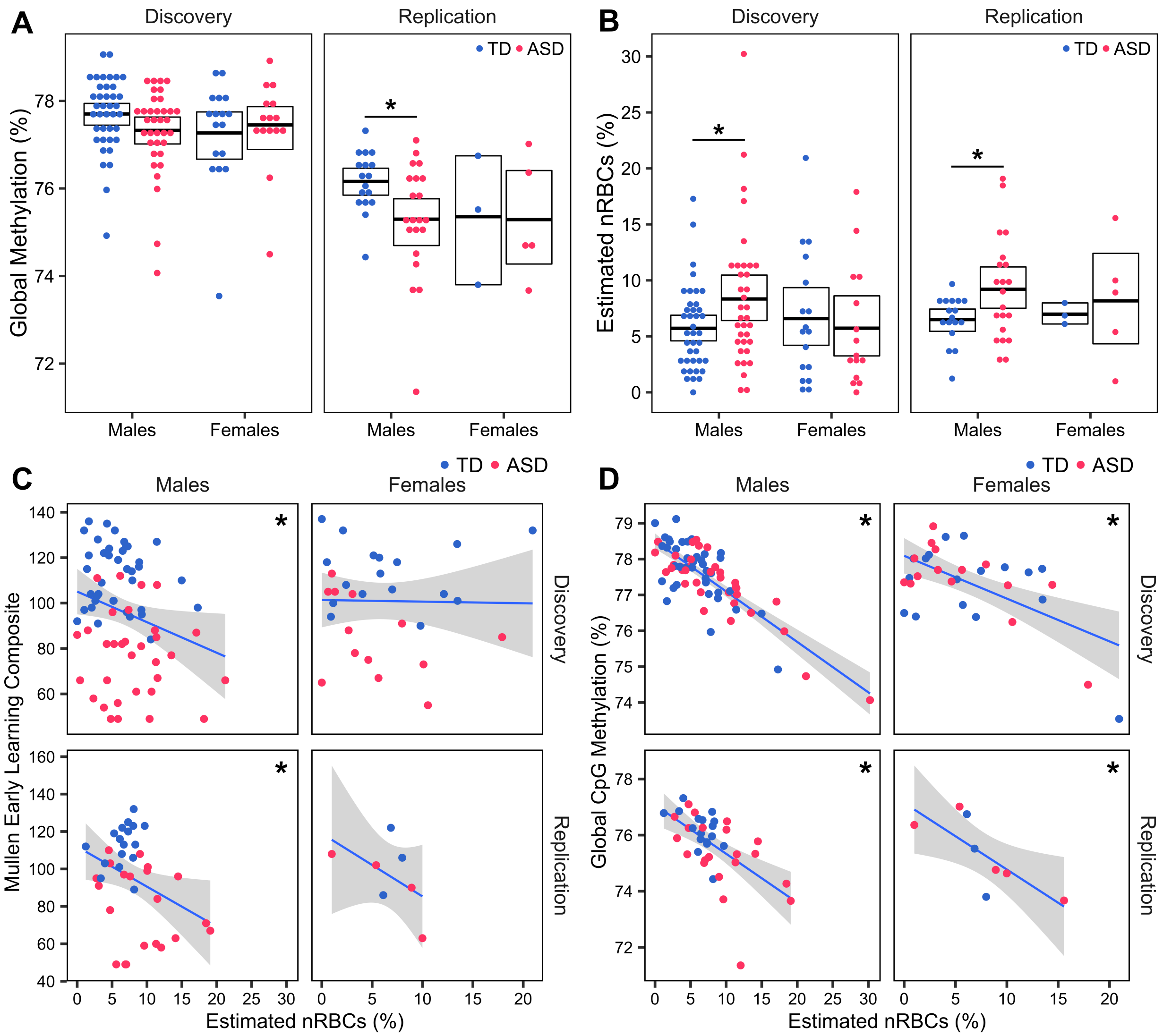


**Supplemental Figure 2. The proportion of nRBCs is associated with behavioral outcome and global CpG methylation in males.** (A) Percent CpG methylation across the genome. Boxes represent mean and 95% confidence limits by nonparametric bootstrapping. Linear model included adjustment for PCR duplicates (pooled males *p* = 0.002, pooled females *p* = 0.70). (B) Estimated proportion of nRBCs. Boxes represent mean and 95% confidence limits by nonparametric bootstrapping (pooled males *p* = 0.003, pooled females *p* = 0.78). (C) Estimated proportion of nRBCs compared to Mullen Early Learning Composite score is plotted by sex and sample set (pooled males *p* = 0.003, pooled females *p* = 0.71). (D) Estimated proportion of nRBCs compared to global CpG methylation is plotted by sex and sample set (pooled males *p* =

1.7E-18, pooled females $p = 3.1E-5$). Significance was tested using linear regression with adjustment for sequencing platform when pooled (* $p < 0.05$; pooled males TD $n = 56$, ASD $n = 56$; pooled females TD $n = 20$, ASD $n = 20$).

**Region-specific differential methylation patterns in umbilical cord blood distinguish males and females later diagnosed with ASD from typically developing controls**

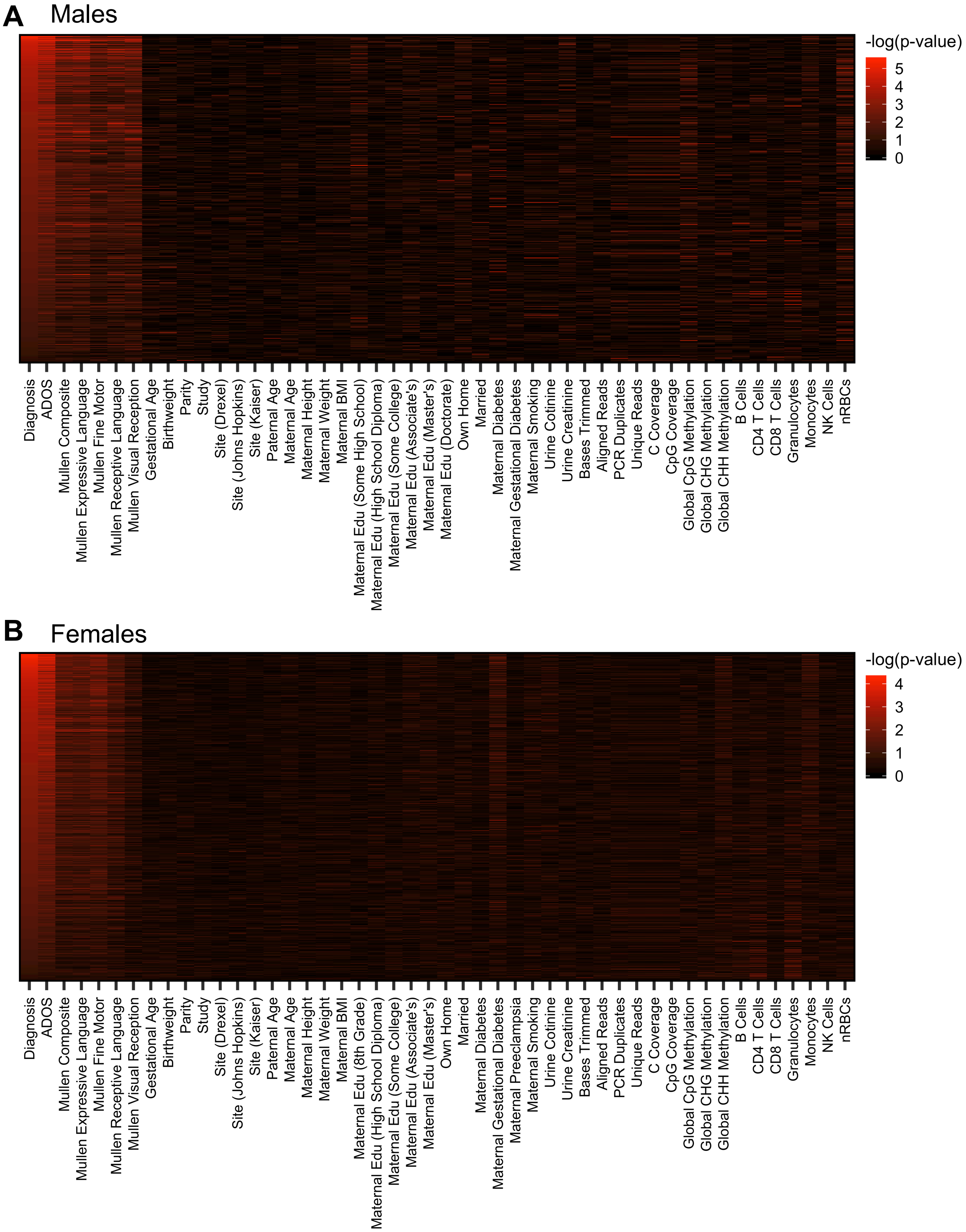

**Supplemental Figure 3. Methylation at ASD DMRs in the discovery set is specifically associated with behavioral outcome.** Raw percent methylation at ASD differentially-methylated regions (DMRs) identified in (A) male or (B) female discovery set subjects was compared with demographic and technical variables. Significance testing was done with linear regression and the -$\log_{10}$(p-value) was plotted (males TD $n$ = 39, ASD $n$ = 35; females TD $n$ = 17, ASD $n$ = 15). Edu, education;

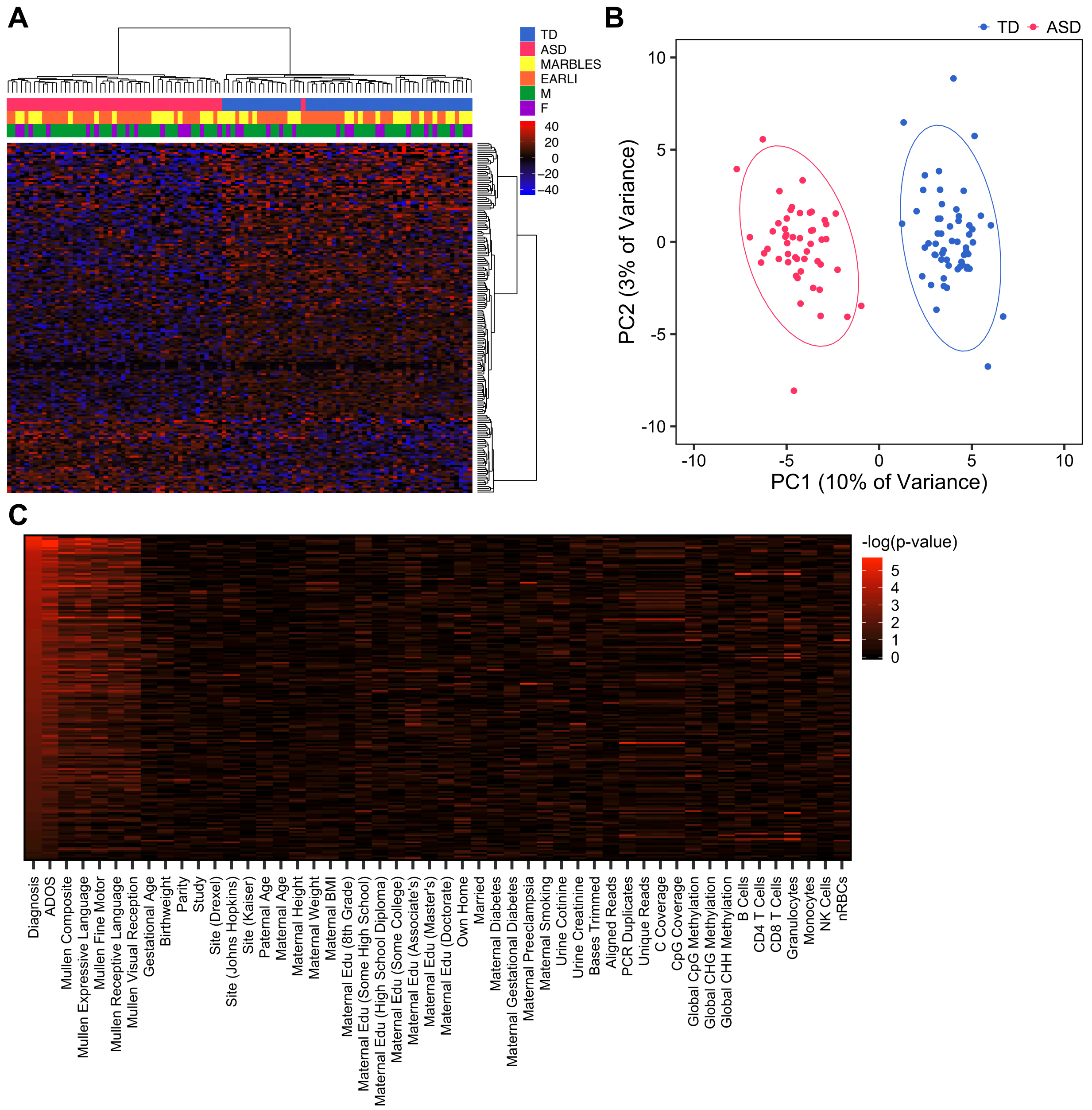

**Supplemental Figure 4. DMRs identified in all discovery subjects distinguish ASD from TD subjects.** (A) Heatmap or (B) principal component analysis (PCA) plot using percent methylation for each sample at ASD DMRs identified in all discovery subjects with adjustment for sex (42 hypermethylated DMRs, 145 hypomethylated DMRs). For heatmap, subjects are colored by diagnostic group and study, and methylation is relative to the mean for each DMR. For PCA plot, ellipses indicate 95% confidence limits. (C) Raw percent methylation at ASD DMRs identified in all discovery subjects was compared with demographic and technical variables. Significance testing was done using linear regression with adjustment for sex and the $-\log_{10}$(p-value) was plotted (males TD $n$ = 39, ASD $n$ = 35; females TD $n$ = 17, ASD $n$ = 15). EARLI, Early Autism Risk Longitudinal Investigation; F, female; M, male; MARBLES, Markers of Autism Risk in Babies - Learning Early Signs;

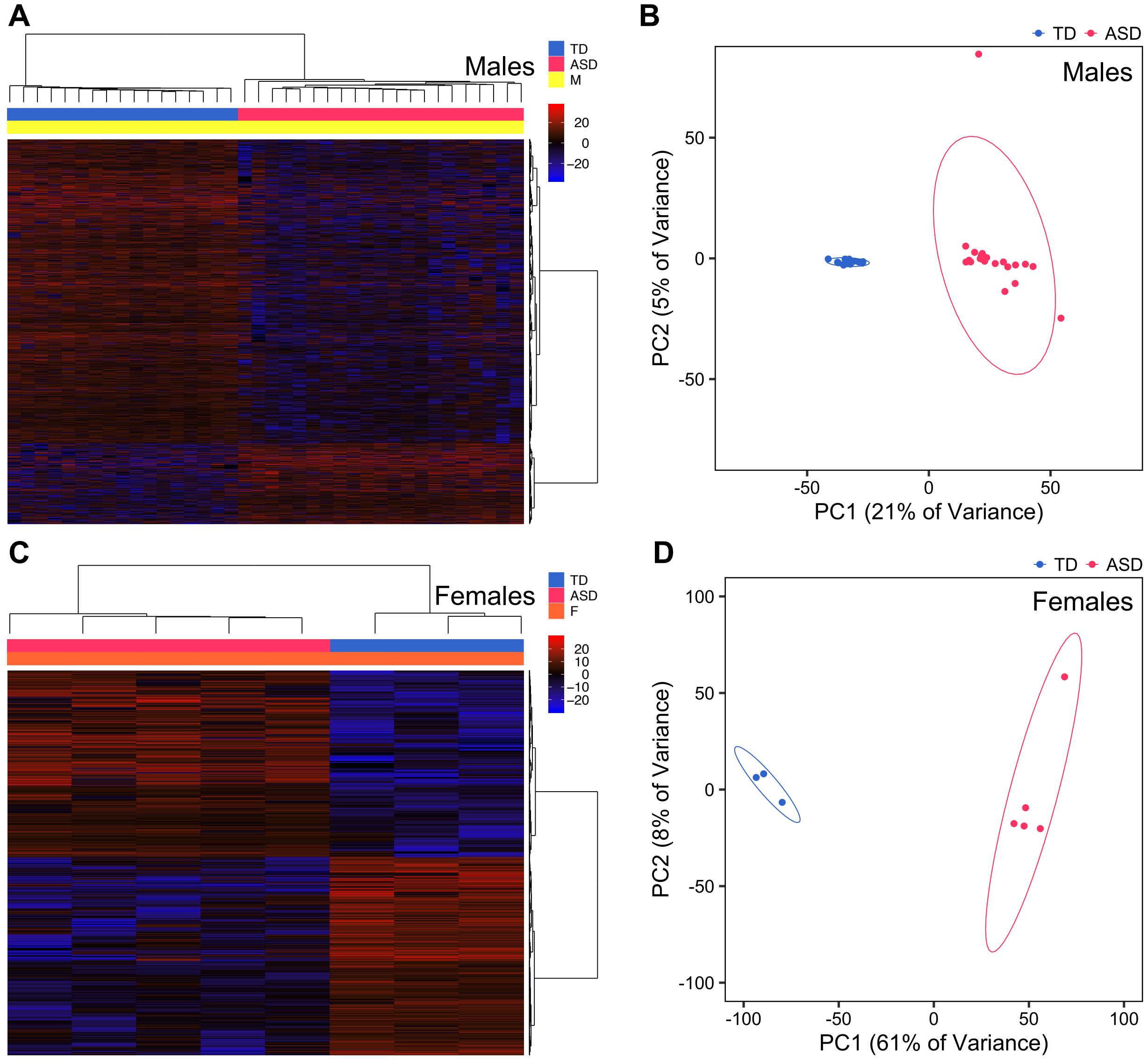


**Supplemental Figure 5. DMRs identified in males and females distinguish ASD from TD subjects in the replication set.** (A) Heatmap or (B) PCA plot using percent methylation for each sample at ASD DMRs identified in male replication subjects (975 hypermethylated DMRs, 3675 hypomethylated DMRs). (C) Heatmap or (D) PCA plot using percent methylation for each sample at ASD DMRs identified in female replication subjects (4232 hypermethylated DMRs, 4496 hypomethylated DMRs). For heatmaps, subjects are colored by diagnostic group and study, and methylation is relative to the mean for each DMR. For PCA plots, ellipses indicate 95% confidence limits (males TD *n* = 17, ASD *n* = 21; females TD *n* = 3, ASD *n* = 5).

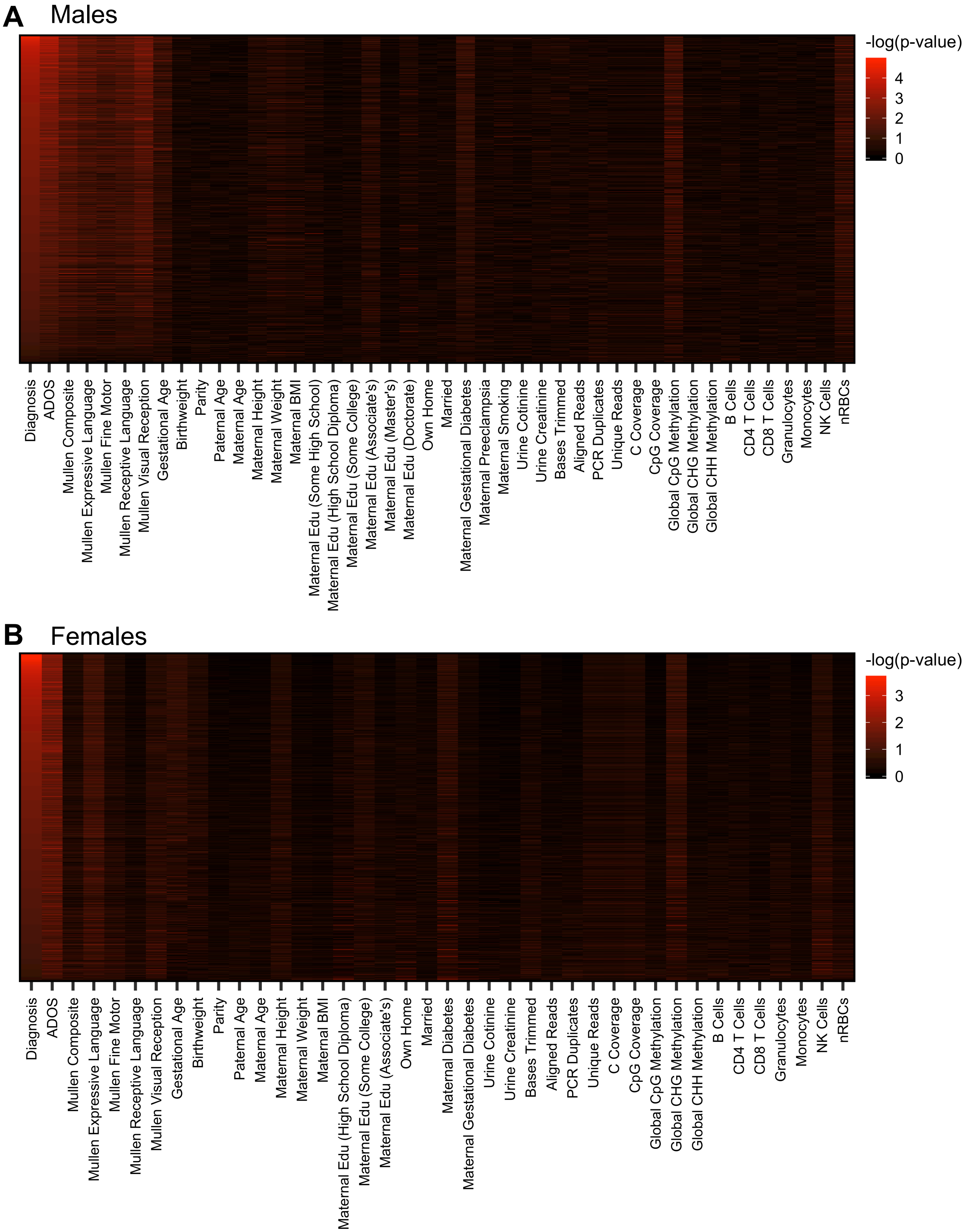
A
Males
-log(p-value)
4
3
2
1
0
Diagnosis
ADOS
Mullen Composite
Mullen Expressive Language
Mullen Fine Motor
Mullen Receptive Language
Mullen Visual Reception
Gestational Age
Birthweight
Parity
Paternal Age
Maternal Age
Maternal Height
Maternal Weight
Maternal BMI
Maternal Edu (Some High School)
Maternal Edu (High School Diploma)
Maternal Edu (Some College)
Maternal Edu (Associate's)
Maternal Edu (Master's)
Maternal Edu (Doctorate)
Own Home
Married
Maternal Gestational Diabetes
Maternal Preeclampsia
Maternal Smoking
Urine Cotinine
Urine Creatinine
Bases Trimmed
Aligned Reads
PCR Duplicates
Unique Reads
C Coverage
CpG Coverage
Global CpG Methylation
Global CHG Methylation
Global CHH Methylation
B Cells
CD4 T Cells
CD8 T Cells
Granulocytes
Monocytes
NK Cells
nRBCs
B
Females
-log(p-value)
3
2
1
0
Diagnosis
ADOS
Mullen Composite
Mullen Expressive Language
Mullen Fine Motor
Mullen Receptive Language
Mullen Visual Reception
Gestational Age
Birthweight
Parity
Paternal Age
Maternal Age
Maternal Height
Maternal Weight
Maternal BMI
Maternal Edu (High School Diploma)
Maternal Edu (Some College)
Maternal Edu (Associate's)
Own Home
Married
Maternal Diabetes
Maternal Gestational Diabetes
Urine Cotinine
Urine Creatinine
Bases Trimmed
Aligned Reads
PCR Duplicates
Unique Reads
C Coverage
CpG Coverage
Global CpG Methylation
Global CHG Methylation
Global CHH Methylation
B Cells
CD4 T Cells
CD8 T Cells
Granulocytes
Monocytes
NK Cells
nRBCs

1

**Supplemental Figure 6. Methylation at ASD DMRs in the replication set is specifically associated with behavioral outcome.** Raw percent methylation at ASD DMRs identified in (A) male or (B) female replication set subjects was compared with demographic and technical variables. Significance testing was done with linear regression and the -$\log_{10}$(p-value) was plotted (males TD $n$ = 17, ASD $n$ = 21; females TD $n$ = 3, ASD $n$ = 5).

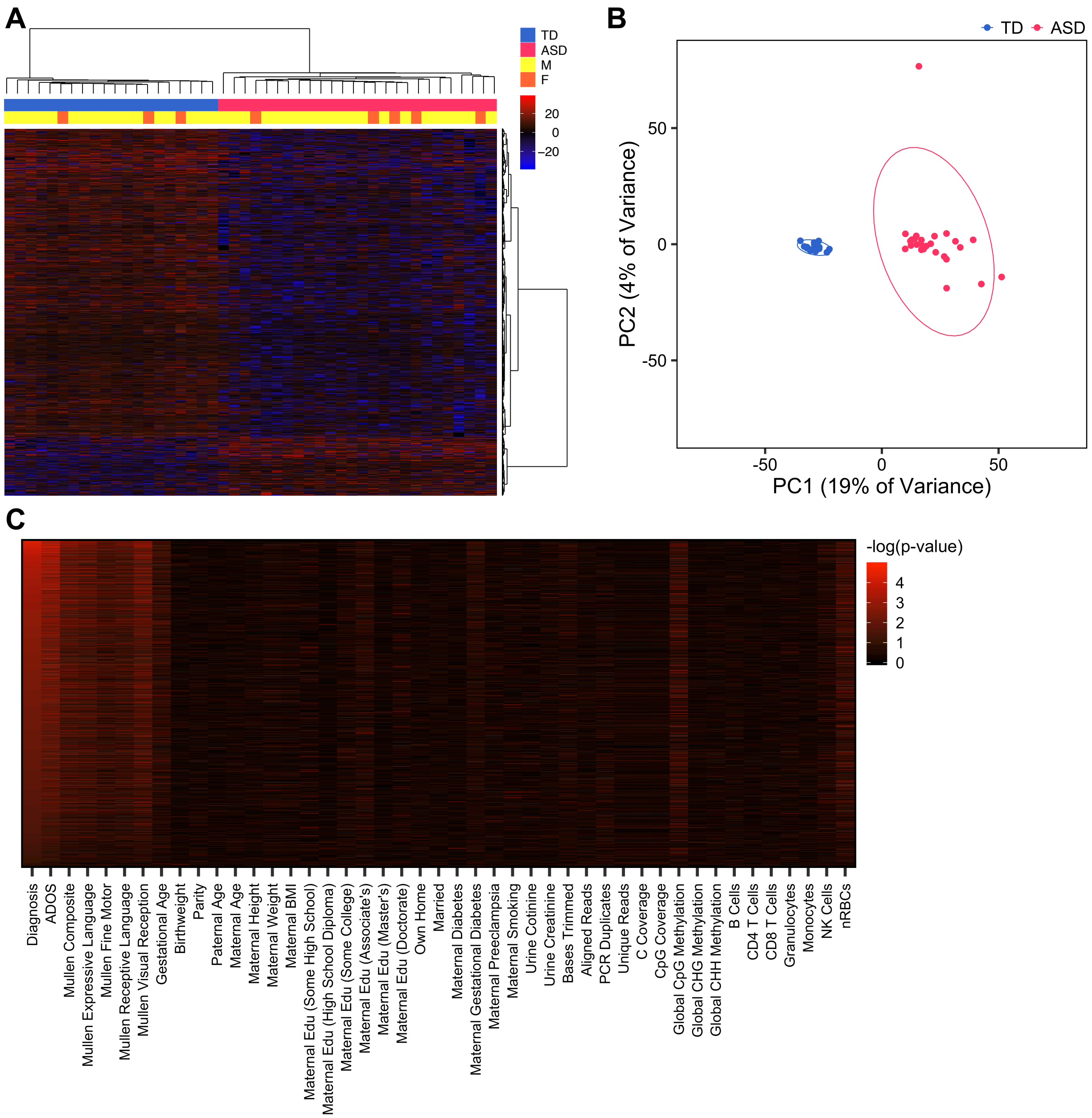


**Supplemental Figure 7. DMRs identified in all replication subjects distinguish ASD from**

**TD subjects.** (A) Heatmap or (B) PCA plot using percent methylation for each sample at ASD DMRs identified in all replication subjects with adjustment for sex (614 hypermethylated DMRs, 3207 hypomethylated DMRs). For heatmap, subjects are colored by diagnostic group and study, and methylation is relative to the mean for each DMR. For PCA plot, ellipses indicate 95% confidence limits. (C) Raw percent methylation at ASD DMRs identified in all replication subjects was compared with demographic and technical variables. Significance testing was done using linear regression with adjustment for sex and the $-\log_{10}$(p-value) was plotted (males TD *n* = 17, ASD *n* = 21; females TD *n* = 3, ASD *n* = 5).

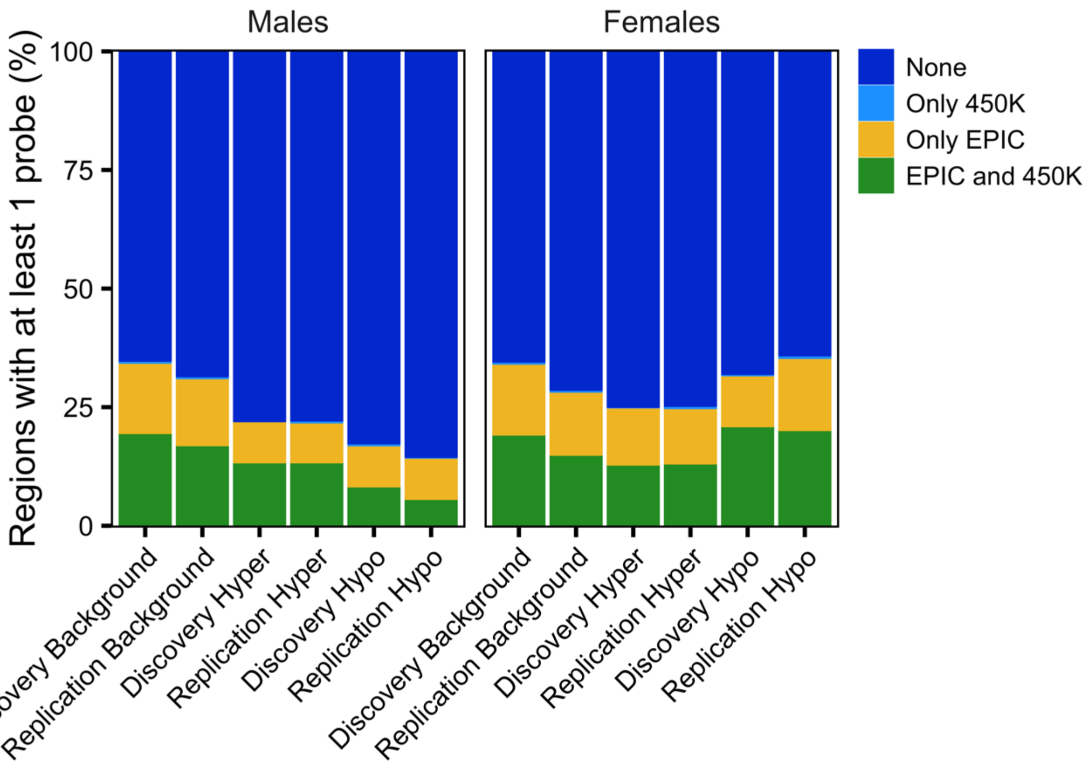


**Supplemental Figure 8. The majority of ASD DMRs do not overlap probes on the 450K and EPIC arrays.** ASD DMRs and background regions in Discovery or Replication Male or Female subjects were overlapped by location with probes on the Infinium HumanMethylation450 (450K) and MethylationEPIC (EPIC) arrays, and the proportion of regions overlapping at least one probe on either or both arrays was plotted (pooled males TD *n* = 56, ASD *n* = 56; pooled females TD *n* = 20, ASD *n* = 20). Hyper, hypermethylated; Hypo, hypomethylated;

**ASD DMRs in umbilical cord blood replicate across independent groups of subjects**

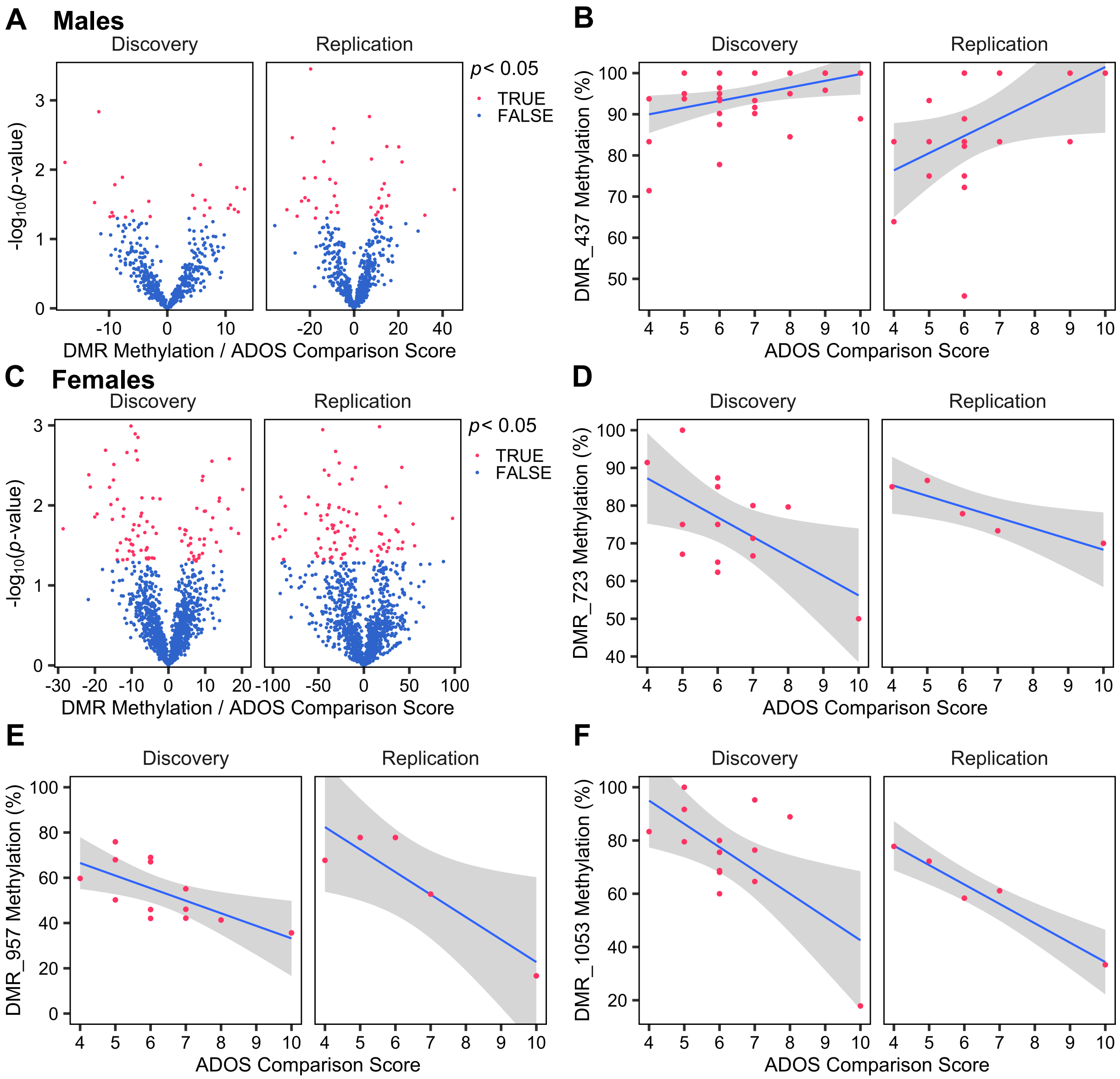


**Supplemental Figure 9. A subset of ASD diagnosis DMRs are associated with ASD severity in independent sample sets.** (A,C) Volcano plots of the association between Discovery DMR methylation and ADOS comparison score for (A) ASD males or (C) ASD females by Discovery or Replication sample set. The x-axis represents the change in percent methylation per unit of the ADOS comparison score. (B,D-F) Scatterplots of ADOS comparison score versus percent methylation at Discovery DMRs with nominal significance ($p < 0.05$) in both Discovery and Replication sample sets in (B) ASD males or (D-F) ASD females (ASD males: Discovery *n* = 35, Replication *n* = 21; ASD females: Discovery *n* = 15, Replication *n* = 5).

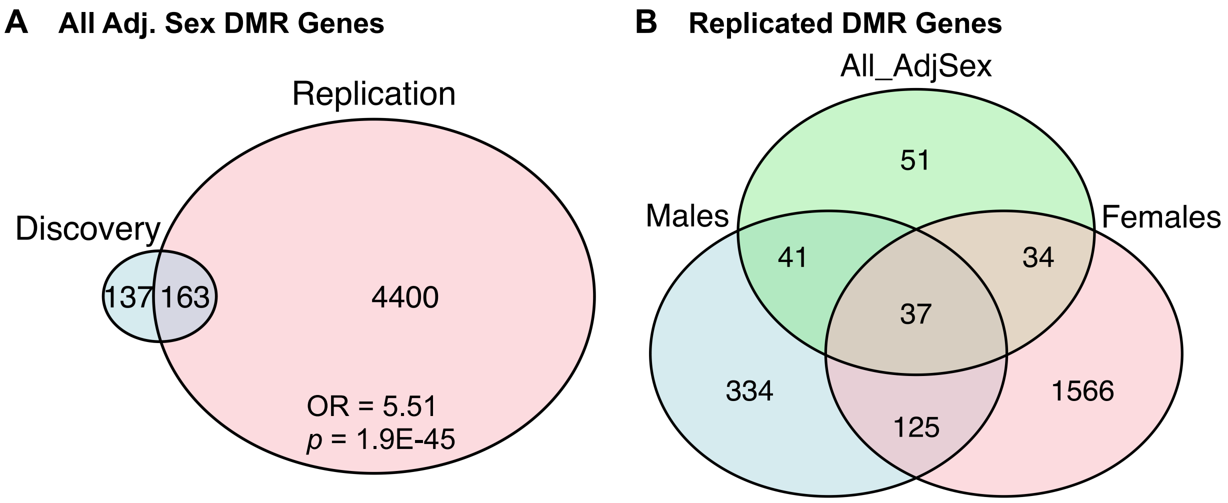


**Supplemental Figure 10. ASD DMRs identified with adjustment for sex miss many genes found when stratifying for sex.** (A) Venn diagram of genes annotated to ASD DMRs with adjustment for sex in all samples in Discovery and Replication sets. Significance was tested using the hypergeometric test and was relative to genes annotated to background regions. (B) Overlap of replicated ASD DMR genes between all samples, males, and females (pooled males TD *n* = 56, ASD *n* = 56; pooled females TD *n* = 20, ASD *n* = 20). Adj, adjusted for; OR, odds ratio;

**<u>Genes in ASD differentially-methylated blocks replicate between independent groups of subjects and are enriched for ASD DMR genes, cadherins, and developmental genes</u>**

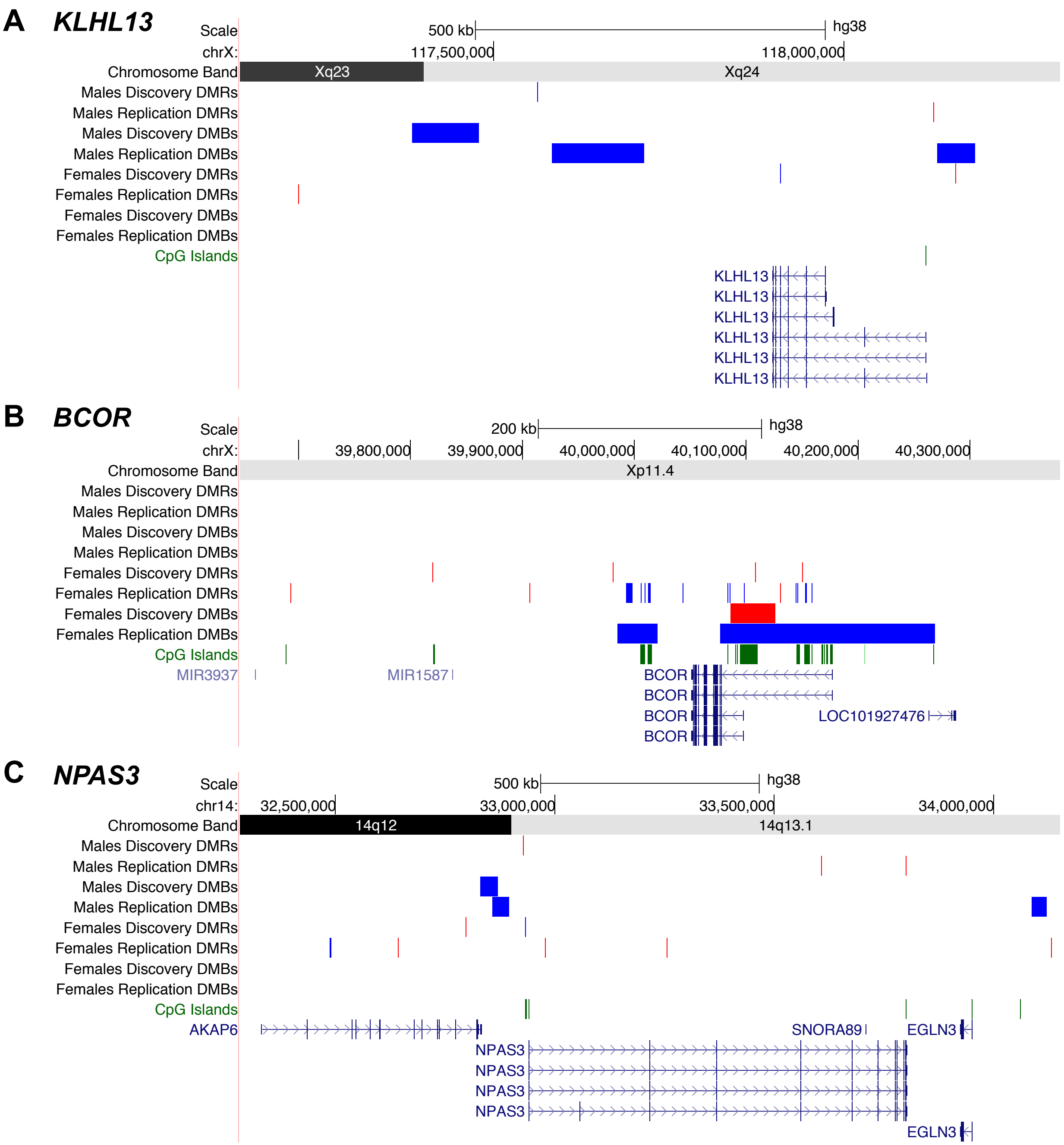


**Supplemental Figure 11. Replicated ASD DMB genes overlap with replicated ASD DMR genes.** Selected regions with replicated DMRs and differentially-methylated blocks (DMBs) in males or females. chr, chromosome;

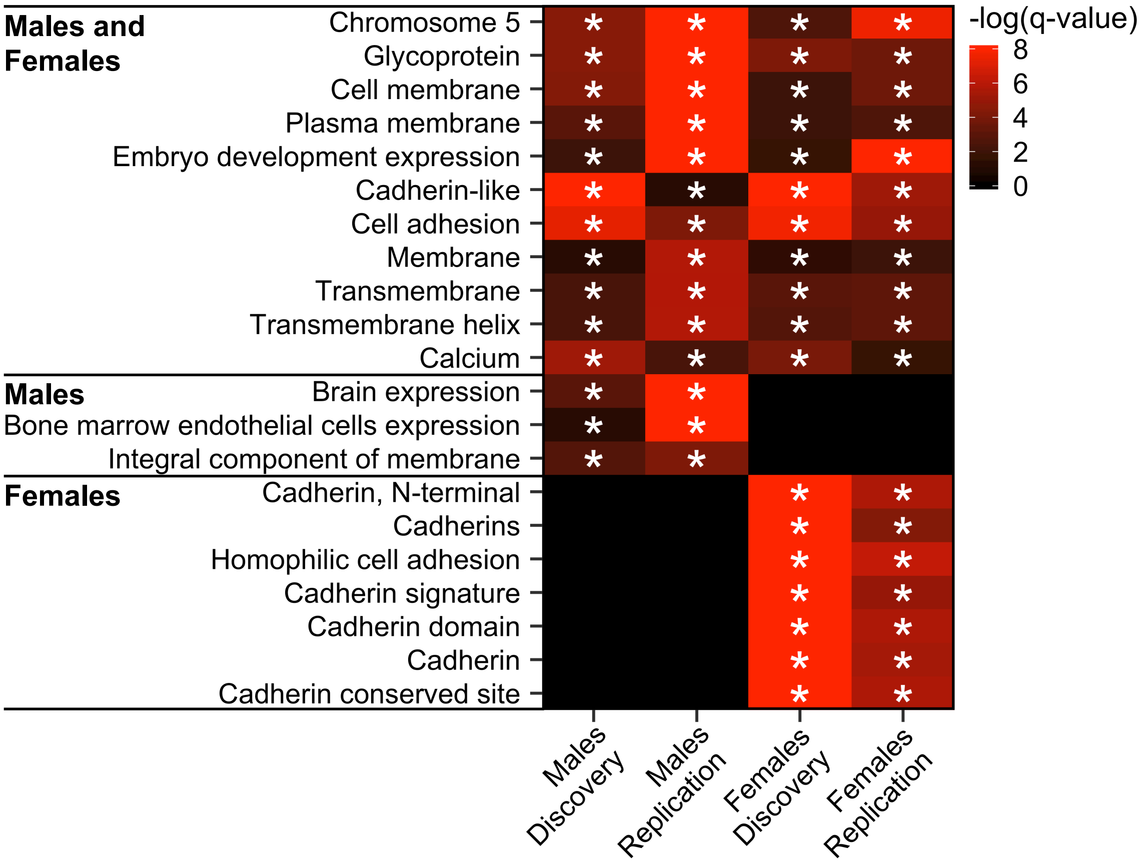


**Supplemental Figure 12. ASD DMB genes are enriched for membrane, cell adhesion, and embryo-expressed genes.** Terms significantly enriched among ASD DMB genes in both discovery and replication sample sets for either males or females (* $q < 0.05$). Heatmaps show -$\log_{10}$($q$-value) for enrichment in genes annotated to DMBs relative to genes annotated to background calculated using the Database for Annotation, Visualization, and Integrated Discovery (DAVID) for all categories. Terms were sorted by replication sex (pooled males TD $n$ = 56, ASD $n$ = 56; pooled females TD $n$ = 20, ASD $n$ = 20).

**<u>Cord blood ASD DMR genes are enriched for neurodevelopmental genes on the X chromosome that are epigenetically dysregulated in ASD brain</u>**

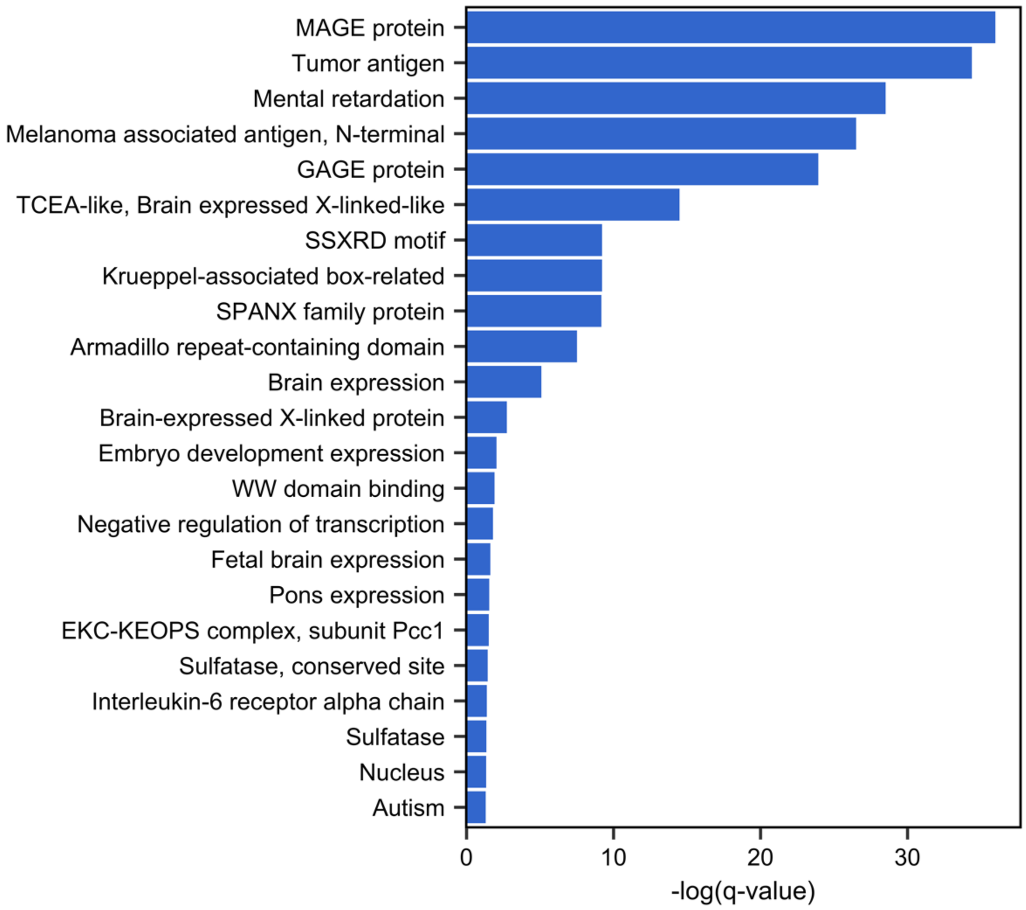


**Supplemental Figure 13. Neurodevelopmental genes are overrepresented on the X chromosome.** Terms significantly enriched among chromosome X genes compared to all genes ($q$ < 0.05). Plot shows -$\log_{10}$($q$-value) for enrichment calculated using DAVID for all categories.

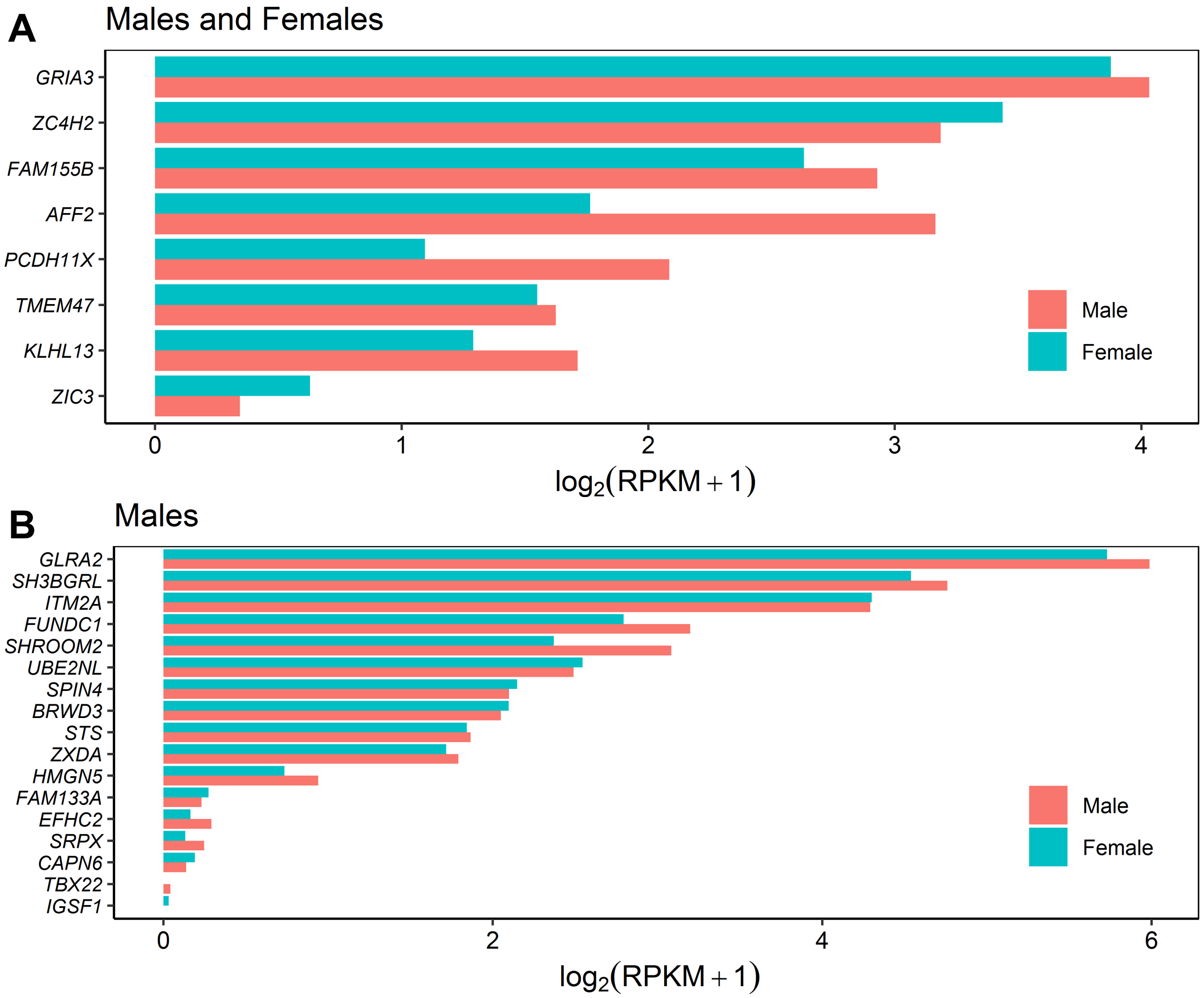


**Supplemental Figure 14. Replicated DMR genes on the X chromosome are expressed in fetal brain.** RNA-seq expression values were obtained from the Allen BrainSpan Atlas of the Developing Human Brain for one male and one female dorsolateral prefrontal cortex at 13 weeks post-conception for X-linked DMR genes replicated in (A) males and females or (B) males only. RPKM, reads per kilobase per million reads;

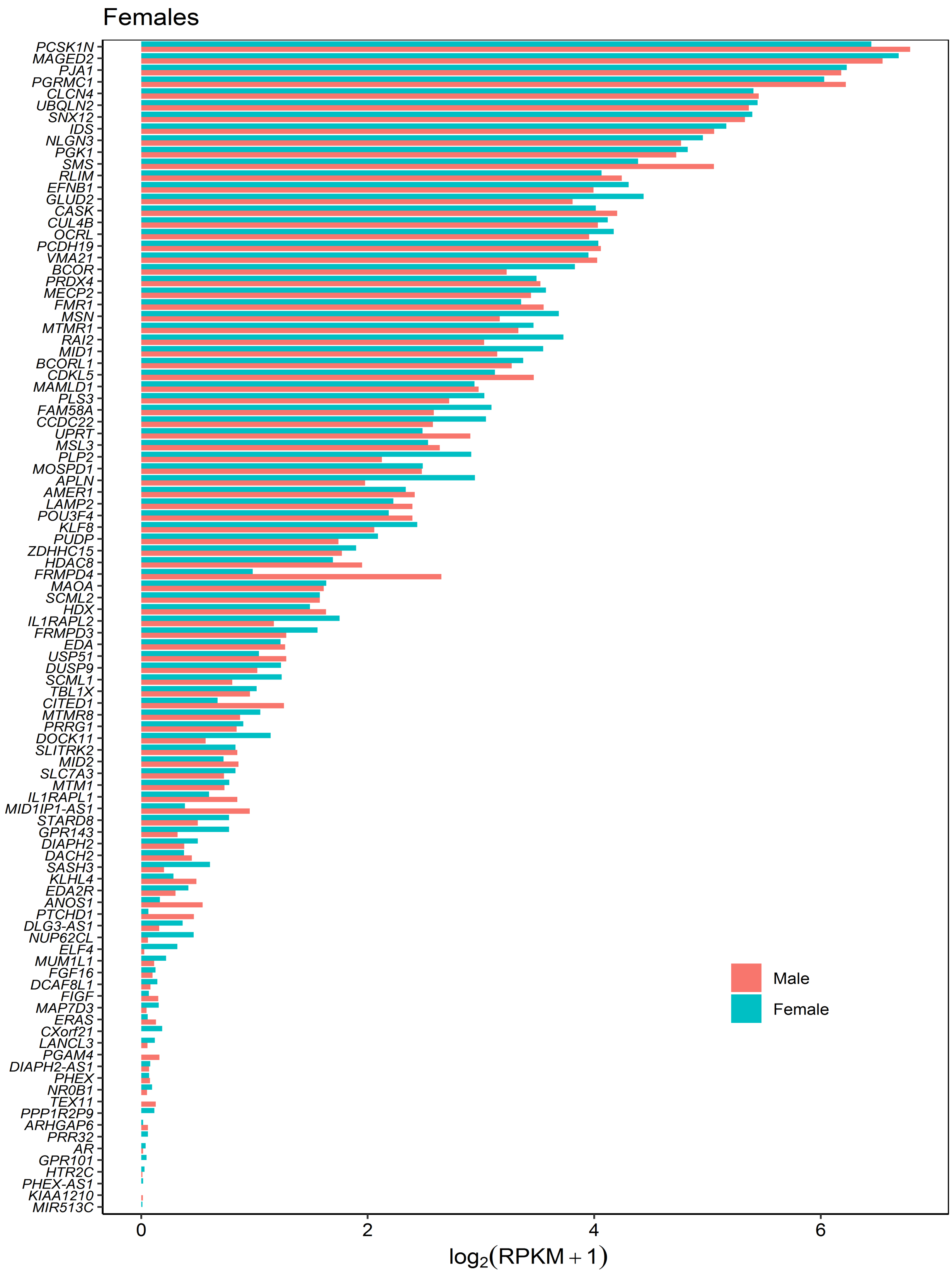
Females
PCSK1N
MAGED2
PJA1
PGRMC1
CLCN4
UBQLN2
SNX12
IDS
NLGN3
PGK1
SMS
RLIM
EFNB1
GLUD2
CASK
CUL4B
OCRL
PCDH19
VMA21
BCOR
PRDX4
MECP2
FMR1
MSN
MTMR1
RAI2
MID1
BCORL1
CDKL5
MAMLD1
PLS3
FAM58A
CCDC22
UPRT
MSL3
PLP2
MOSPD1
APLN
AMER1
LAMP2
POU3F4
KLF8
PUDP
ZDHHC15
HDAC8
FRMPD4
MAOA
SCML2
HDX
IL1RAPL2
FRMPD3
EDA
USP51
DUSP9
SCML1
TBL1X
CITED1
MTMR8
PRRG1
DOCK11
SLITRK2
MID2
SLC7A3
MTM1
IL1RAPL1
MID1IP1-AS1
STARD8
GPR143
DIAPH2
DACH2
SASH3
KLHL4
EDA2R
ANOS1
PTCHD1
DLG3-AS1
NUP62CL
ELF4
MUM1L1
FGF16
DCAF8L1
FIGF
MAP7D3
ERAS
CXorf21
LANCL3
PGAM4
DIAPH2-AS1
PHEX
NR0B1
TEX11
PPP1R2P9
ARHGAP6
PRR32
AR
GPR101
HTR2C
PHEX-AS1
KIAA1210
MIR513C
0
2
4
6
log2(RPKM + 1)
Male
Female

**Supplemental Figure 15. Female-specific replicated DMR genes on the X chromosome are expressed in fetal brain.** RNA-seq expression values were obtained from the Allen BrainSpan Atlas of the Developing Human Brain for one male and one female dorsolateral prefrontal cortex at 13 weeks post-conception for X-linked DMR genes replicated in females only.

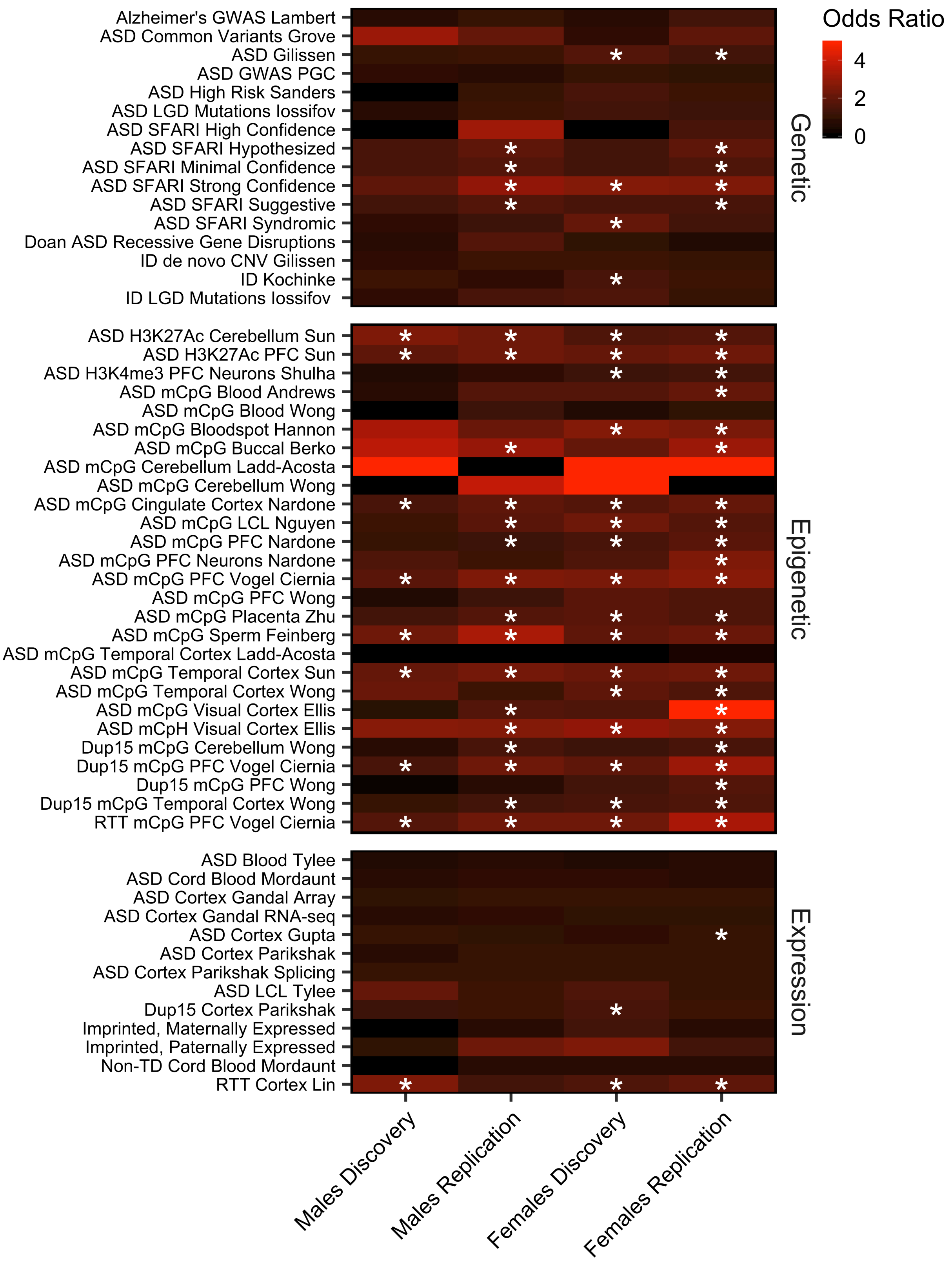
Odds Ratio
4
2
0
Genetic
Epigenetic
Expression
Alzheimer's GWAS Lambert
ASD Common Variants Grove
ASD Gilissen
ASD GWAS PGC
ASD High Risk Sanders
ASD LGD Mutations Iossifov
ASD SFARI High Confidence
ASD SFARI Hypothesized
ASD SFARI Minimal Confidence
ASD SFARI Strong Confidence
ASD SFARI Suggestive
ASD SFARI Syndromic
Doan ASD Recessive Gene Disruptions
ID de novo CNV Gilissen
ID Kochinke
ID LGD Mutations Iossifov
ASD H3K27Ac Cerebellum Sun
ASD H3K27Ac PFC Sun
ASD H3K4me3 PFC Neurons Shulha
ASD mCpG Blood Andrews
ASD mCpG Blood Wong
ASD mCpG Bloodspot Hannon
ASD mCpG Buccal Berko
ASD mCpG Cerebellum Ladd-Acosta
ASD mCpG Cerebellum Wong
ASD mCpG Cingulate Cortex Nardone
ASD mCpG LCL Nguyen
ASD mCpG PFC Nardone
ASD mCpG PFC Neurons Nardone
ASD mCpG PFC Vogel Ciernia
ASD mCpG PFC Wong
ASD mCpG Placenta Zhu
ASD mCpG Sperm Feinberg
ASD mCpG Temporal Cortex Ladd-Acosta
ASD mCpG Temporal Cortex Sun
ASD mCpG Temporal Cortex Wong
ASD mCpG Visual Cortex Ellis
ASD mCpH Visual Cortex Ellis
Dup15 mCpG Cerebellum Wong
Dup15 mCpG PFC Vogel Ciernia
Dup15 mCpG PFC Wong
Dup15 mCpG Temporal Cortex Wong
RTT mCpG PFC Vogel Ciernia
ASD Blood Tylee
ASD Cord Blood Mordaunt
ASD Cortex Gandal Array
ASD Cortex Gandal RNA-seq
ASD Cortex Gupta
ASD Cortex Parikshak
ASD Cortex Parikshak Splicing
ASD LCL Tylee
Dup15 Cortex Parikshak
Imprinted, Maternally Expressed
Imprinted, Paternally Expressed
Non-TD Cord Blood Mordaunt
RTT Cortex Lin
Males Discovery
Males Replication
Females Discovery
Females Replication

**Supplemental Figure 16. Cord blood ASD DMR genes are significantly enriched for epigenetically dysregulated genes in ASD brain.** All gene sets overlapped with ASD DMR genes in males or females (* $q < 0.05$). Heatmaps show odds ratios for enrichment in genes annotated to DMRs relative to genes annotated to background calculated with Fisher's exact test for previously published studies of ASD and other neurological disorders. P-values were adjusted using the FDR method for the total number of gene lists compared (pooled males TD $n = 56$, ASD $n = 56$; pooled females TD $n = 20$, ASD $n = 20$). Ac, acetylation; Dup15, Chromosome 15q11-q13 Duplication syndrome; GWAS, genome wide association study; H3K27, histone 3 lysine 27; H3K4, histone 3 lysine 4; ID, intellectual disability; LCL, lymphoblastoid cell line; LGD, likely gene disrupting; mCpG, CpG methylation; mCpH, CpH methylation; me3, trimethylation; Non-TD, non-typically developing; PFC, prefrontal cortex; PGC, Psychiatric Genomics Consortium; RTT, Rett syndrome; SFARI, Simons Foundation Autism Research Initiative;

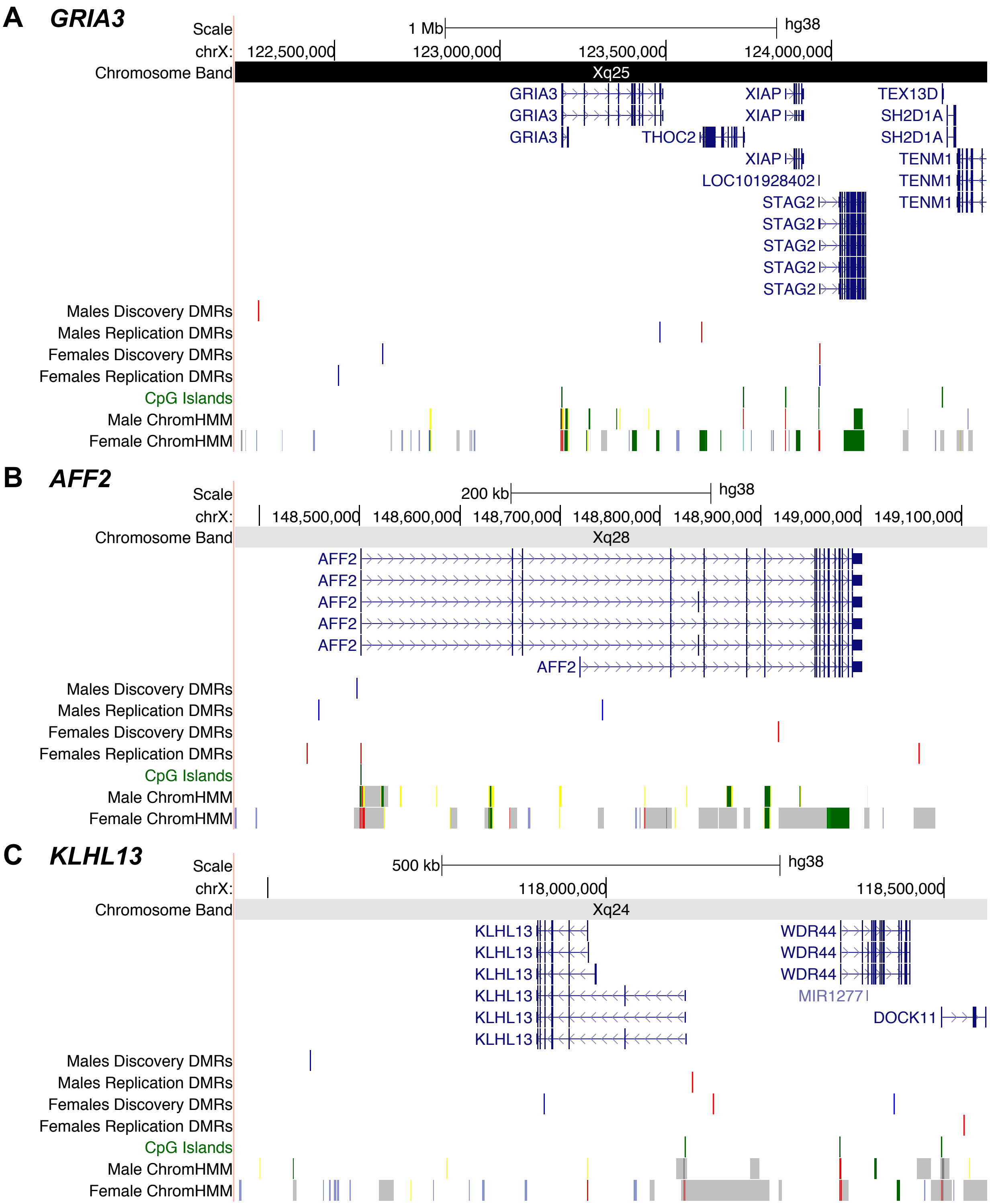


**Supplemental Figure 17. Selected regions with replicated sex-independent DMR genes on the X chromosome.** All shown regions replicated in males and females, were reported in a previous epigenetic study of ASD, and are expressed in fetal brain. Fetal brain ChromHMM chromatin state tracks were obtained from the Roadmap Epigenomics Project.

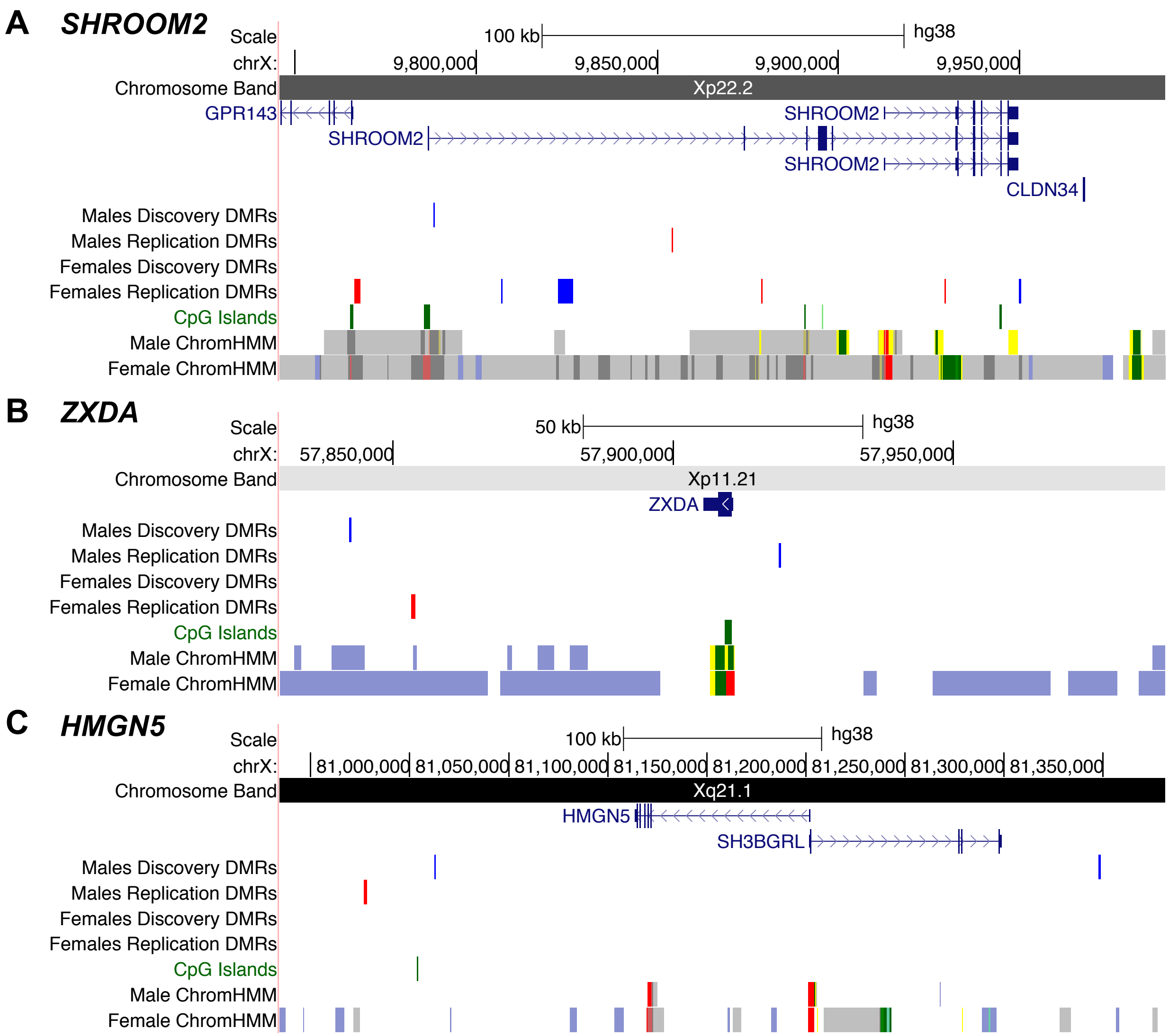


**Supplemental Figure 18. Selected regions with replicated male-specific DMR genes on the X chromosome.** All shown regions replicated in males only, were reported in a previous epigenetic study of ASD, and are expressed in fetal brain. Fetal brain ChromHMM chromatin state tracks were obtained from the Roadmap Epigenomics Project.

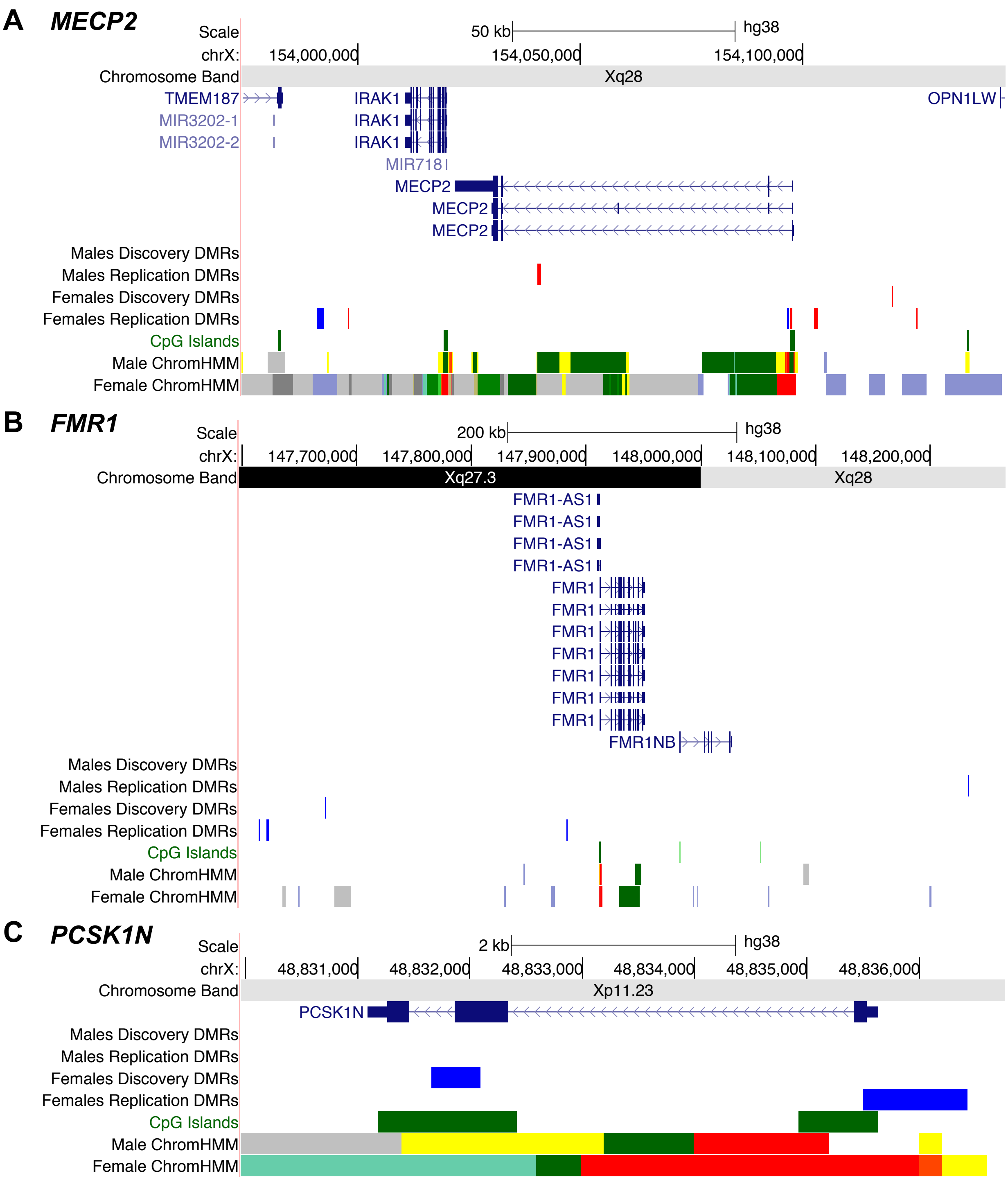


**Supplemental Figure 19. Selected regions with replicated female-specific DMR genes on the X chromosome.** All shown regions replicated in females only, were reported in a previous epigenetic study of ASD, and are expressed in fetal brain. Fetal brain ChromHMM chromatin state tracks were obtained from the Roadmap Epigenomics Project.

**<u>ASD DMRs are enriched for a pan-tissue epigenomic signature that differs between males and females on the X chromosome</u>**

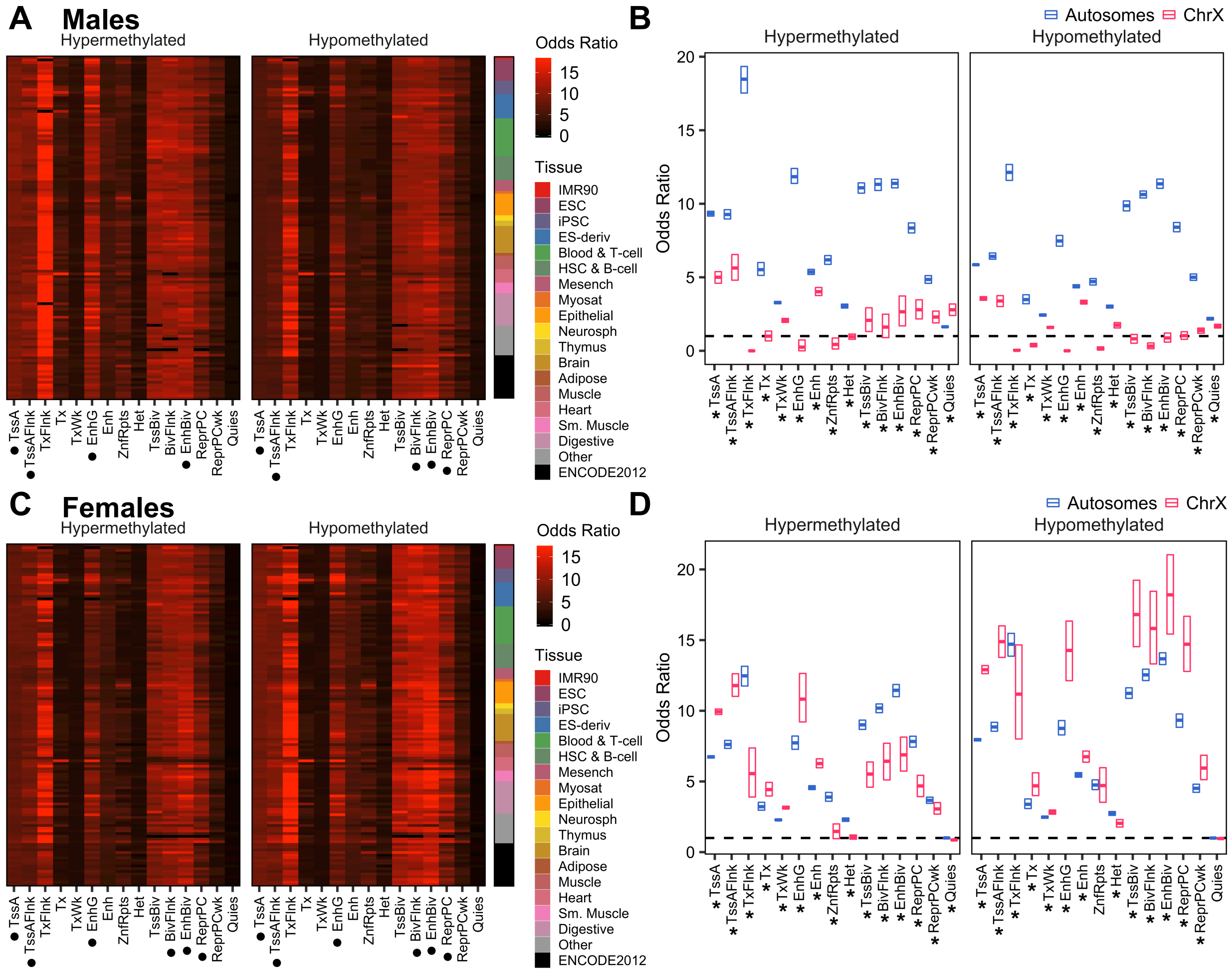


**Supplemental Figure 20. ASD DMRs in replication subjects are differentially enriched for chromatin states on the X chromosome.** ASD DMRs were overlapped with 15-state model ChromHMM segmentations from 127 cell types in the Roadmap Epigenomics Project using the Locus Overlap Analysis (LOLA) R package. (A,C) The enrichment odds ratio was plotted for hypermethylated and hypomethylated DMRs identified in (A) males or (C) females from the replication set. Top enriched (•) chromatin states were identified as those with odds ratio and -log(*q*-value) of at least the median value for that DMR set and with $q < 0.05$ for more than half of all cell types. (B,D) The enrichment odds ratio was plotted for hypermethylated and hypomethylated DMRs on autosomes or the X chromosome identified in (B) males or (D) females from the replication set. Boxes represent mean and 95% confidence limits by nonparametric bootstrapping. Significance of differential enrichment of X chromosome compared to autosome DMRs was assessed by paired t-test of odds ratios for each cell type. P-values were adjusted for the number of chromatin states using the FDR method (* $q < 0.05$, males TD $n = 17$, ASD $n = 21$; females TD $n = 3$, ASD $n = 5$). BivFlnk, flanking bivalent transcription start site or enhancer; Enh, enhancer; EnhBiv, bivalent enhancer; EnhG, genic enhancer; Het, heterochromatin; Quies, quiescent region; ReprPC, polycomb-repressed region; ReprPCWk, weak polycomb-repressed region; TssA, active transcription start site; TssAFlnk, flanking active transcription start site; TssBiv, bivalent transcription start site; Tx, strong transcription; TxFlnk, transcribed at gene 5’ and 3’; TxWk, weak transcription; ZnfRpts, zinc finger genes and repeats;

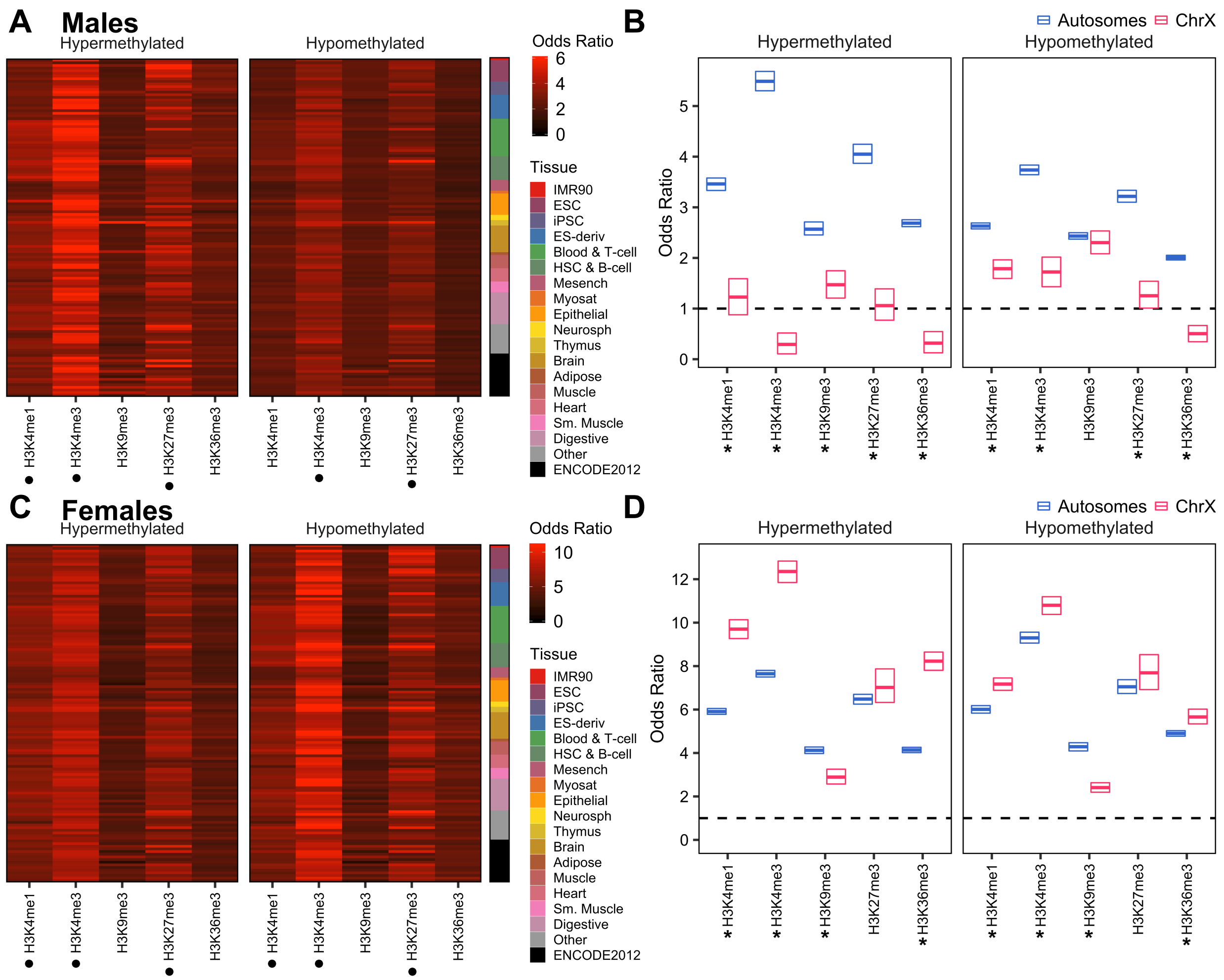


**Supplemental Figure 21. ASD DMRs in discovery subjects are differentially enriched for histone PTMs on the X chromosome.** ASD DMRs were overlapped with histone post-translational modification (PTM) peaks from 127 cell types in the Roadmap Epigenomics Project using LOLA. (A,C) The enrichment odds ratio was plotted for hypermethylated and hypomethylated DMRs identified in (A) males or (C) females from the discovery set. Top enriched (•) histone PTMs were identified as those with odds ratio and -log(*q*-value) of at least the median value for that DMR set and with $q < 0.05$ for more than half of all cell types. (B,D) The enrichment odds ratio was plotted for hypermethylated and hypomethylated DMRs on autosomes or the X chromosome identified in (B) males or (D) females from the discovery set. Boxes represent mean and 95% confidence limits by nonparametric bootstrapping. Significance of differential enrichment of X chromosome compared to autosome DMRs was assessed by paired t-test of odds ratios for each cell type. P-values were adjusted for the number of histone PTMs using the FDR method (* $q < 0.05$, males TD $n = 39$, ASD $n = 35$; females TD $n = 17$, ASD $n = 15$). me1, monomethylation; H3K9, histone 3 lysine 9; H3K36, histone 3 lysine 36;

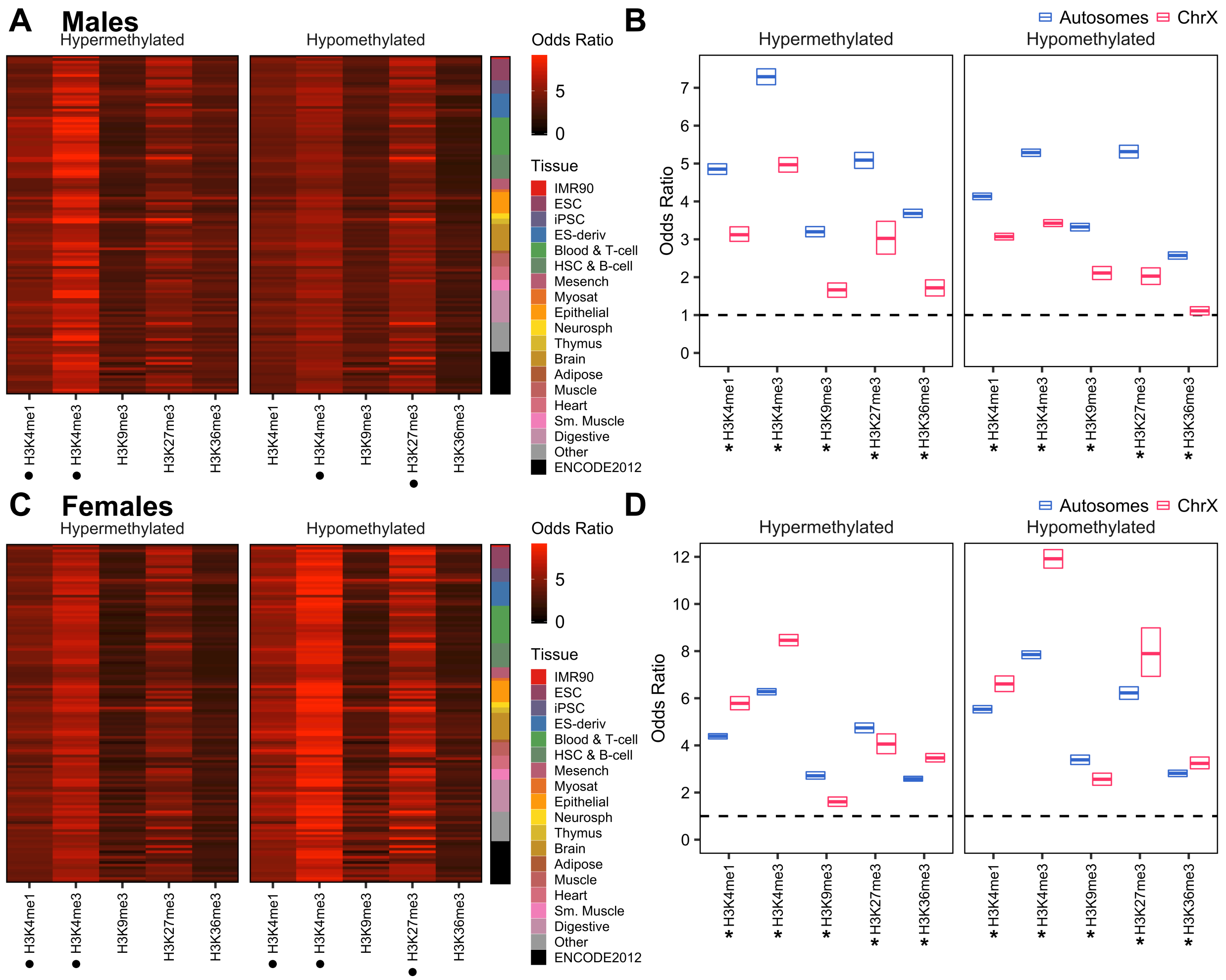


**Supplemental Figure 22. ASD DMRs in replication subjects are differentially enriched for histone PTMs on the X chromosome.** ASD DMRs were overlapped with histone PTM peaks from 127 cell types in the Roadmap Epigenomics Project using LOLA. (A,C) The enrichment odds ratio was plotted for hypermethylated and hypomethylated DMRs identified in (A) males or (C) females from the replication set. Top enriched (•) histone PTMs were identified as those with odds ratio and -log($q$-value) of at least the median value for that DMR set and with $q < 0.05$ for more than half of all cell types. (B,D) The enrichment odds ratio was plotted for hypermethylated and hypomethylated DMRs on autosomes or the X chromosome identified in (B) males or (D) females from the replication set. Boxes represent mean and 95% confidence limits by nonparametric bootstrapping. Significance of differential enrichment of X chromosome compared to autosome DMRs was assessed by paired t-test of odds ratios for each cell type. P-values were adjusted for the number of histone PTMs using the FDR method (* $q < 0.05$, males TD $n = 17$, ASD $n = 21$; females TD $n = 3$, ASD $n = 5$).

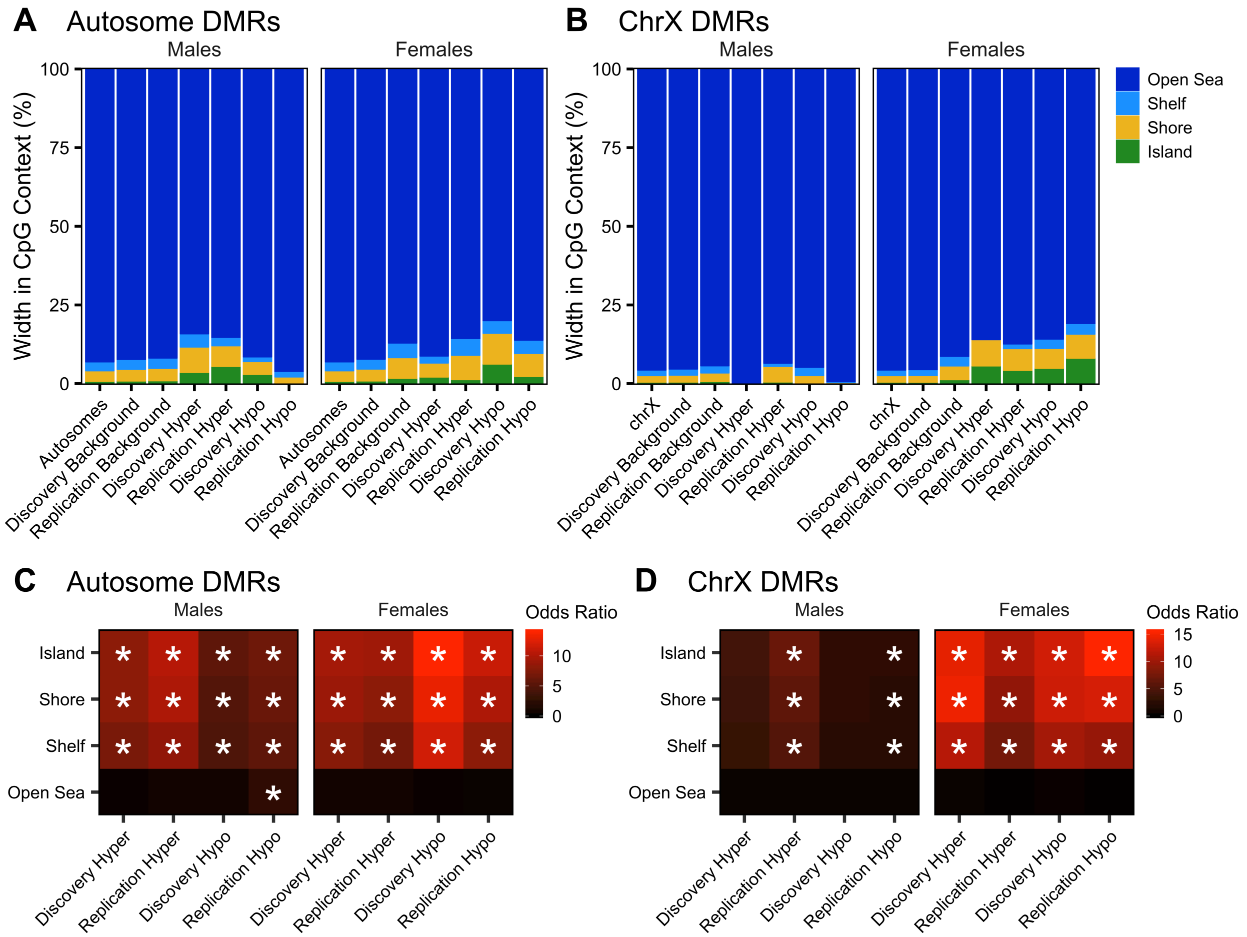


**Supplemental Figure 23. ASD DMRs on the X chromosome are enriched near CpG islands only in females.** ASD DMRs and background regions on (A) autosomes or (B) chrX were intersected with CpG islands, shores, shelves, and open sea as defined in the annotatr R package, and the proportion of total basepairs in each of these contexts was plotted. ASD

DMRs on (C) autosomes or (D) chrX were overlapped with CpG contexts using LOLA and the enrichment odds ratio relative to background regions was plotted (* $q < 0.05$, pooled males TD $n = 56$, ASD $n = 56$; pooled females TD $n = 20$, ASD $n = 20$).
